# Pervasive cooperative mutational effects on multiple catalytic enzyme traits emerge via long-range conformational dynamics

**DOI:** 10.1101/2020.04.14.041590

**Authors:** Carlos G. Acevedo-Rocha, Aitao Li, Lorenzo D’Amore, Sabrina Hoebenreich, Joaquin Sanchis, Paul Lubrano, Matteo P. Ferla, Marc Garcia-Borràs, Sílvia Osuna, Manfred T. Reetz

## Abstract

Multidimensional fitness landscapes provide insights into the molecular basis of laboratory and natural evolution. Yet such efforts are rare and focus only on limited protein families and a single enzyme trait, with little concern about the relationship between protein epistasis and conformational dynamics. Here, we report the first multiparametric fitness landscape for a cytochrome P450 monooxygenase that was engineered for the regio- and stereoselective hydroxylation of a steroid. We developed a computational program to automatically quantify non-additive effects among all possible mutational pathways, finding pervasive cooperative sign and magnitude epistasis on multiple catalytic traits. By using quantum mechanics and molecular dynamics simulations, we show that these effects are modulated by long-range interactions in loops, helices and beta-strands that gate the substrate access channel allowing for optimal catalysis. Our work highlights the importance of conformational dynamics on epistasis in an enzyme involved in secondary metabolism and offers lessons for engineering P450s.

## Introduction

Directed evolution constitutes a powerful tool for optimizing protein properties including activity, substrate scope, selectivity, stability, allostery or binding affinity. By applying iterative rounds of gene mutagenesis, expression and screening (or selection), proteins have been engineered for developing more efficient industrial biocatalytic processes^1–4^. Directed evolution has also provided important insights into the relationship between protein sequence and function^4–6^, yet understanding the intricacies of non-additive epistatic effects remains a challenge^7^. Epistasis means that the phenotypic consequences of a mutation depend on the genetic background^8–11^. Epistatic effects can be negative (antagonistic/deleterious) or positive (synergistic/cooperative) if the respective predictive value is smaller or greater in sign/magnitude than the expected value under additivity. Based on studies of natural and laboratory protein evolution, negative^10^ or positive^11^ epistasis is more widespread than originally thought^7^. Importantly, positive epistasis increases the evolution of new protein functions because it allows access to mutational pathways that avoid deleterious downfalls. For fundamental and practical reasons, it is thus important to determine the existence, type and molecular basis of epistasis in protein evolution. Epistatic effects can arise between residues that are located closely or away from each other via long-range indirect interactions, both mechanisms involving sometimes direct or indirect substrate binding^11^. These global epistatic effects may be mediated by changes in the protein conformational dynamics.

Proteins have the inherent ability to adopt a variety of thermally accessible conformational states, which play a key role in protein evolvability and activity^12,13^. Along the catalytic cycle, enzymes can adopt multiple conformations important for substrate binding or product release^14,15^, and conformational change can be rate-limiting in some cases^16,17^. Much debated is the existence of a link between active site dynamics and the chemical step^18,19^ Some studies have suggested that mutations remote from the enzyme active site may directly impact the energetically accessible conformational states, thereby influencing catalysis^20–22^. This has been shown by means of crystal structures and NMR spectra of mutants along evolutionary pathways^20,23,24^ together with computational assistance^25–29^. Molecular Dynamics (MD) simulations allow the reconstruction of the enzyme conformational landscape, and how this is altered by mutations introduced by laboratory evolution^28–30^. Tuning the enzyme conformational dynamics has been recently identified as key for novel activity^21,24,29–31^.

Understanding the connection between conformational dynamics and epistasis has been limited to a few model enzymes (mainly beta-lactamases) and a single protein trait (usually activity) as a measure of fitness (this term originally refers to the reproductive success of organisms but it can be applied to protein activity, selectivity or stability)^32–35^. This contrasts with directed evolution where often two or more traits (e.g., activity and selectivity or stability) are sought for practical purposes^4,36^. Therefore, connecting epistasis to conformational dynamics increases our understanding of proteins. In turn, analysing non-additive epistatic effects can be expected to benefit *in silico* directed evolution^37^.

In the present work, we used a combination of enzyme kinetics and computational approaches to investigate epistatic effects and conformational dynamics in the stepwise evolution of a cytochrome P450 monooxygenase (CYP) engineered for the highly active, regio- and stereoselective oxidative hydroxylation of a steroid as a non-natural substrate^38^. To determine epistatic effects effectively, we developed a computational program that can be applied to any protein and catalytic trait (https://epistasis.mutanalyst.com/). Unexpectedly, we found pervasive positive epistatic effects on multiple catalytic traits, with selectivity and activity being generally characterized by sign and magnitude epistasis. We found that the analysis of the link between protein epistasis and conformational dynamics reveals the increasing optimization of activity and selectivity along all evolutionary trajectories through fine tuning of loops, helices and beta-strands that gate active site entrance and modulate the active site by long-range networks of interactions. Our study offers guiding principles for the simultaneous engineering of both activity and selectivity in a model CYP member.

## Results

### Multiple parameters define the biocatalytic landscape in P450_BM3_

The haem-containing CYP super family is involved in the biosynthesis and degradation of organic molecules in a wide range of secondary metabolic pathways^39^ and thus have many applications in biocatalysis^40,41^. Previously, we achieved the stereo- and regioselective hydroxylation of testosterone (**1**) by evolving the self-sufficient *Bacillus megaterium* cytochrome P450_BM3_ monooxygenase^38^. P450_BM3_ is one of the most active and versatile CYPs that oxidizes long fatty acids as the natural substrates, and there are various 3D structures of the haem domain alone without the reductase domain^41^. However, wild type (WT) does not accept steroids, which is the reason why we chose mutant F87A as the starting enzyme. While F87A accepts **1**, it provides in a whole cell system only ∼20% conversion with formation of a 1:1 mixture of 2β-hydroxytestosterone (**2**) and 15β-hydroxytestosterone (**3**)^38^. Combinatorial saturation mutagenesis at the randomization site R47/T49/Y51 allowed the evolution of mutant R47I/T49I/Y51I/F87A (III) displaying 94% 2β-selectivity and 68% conversion of **1** in whole cell reactions^38^ (**Fig. 1a**). The mechanism is known to involve a radical process in which the catalytically active haem-Fe=O (Cpd I) abstracts an H-atom from aliphatic C-H followed by a fast C-O forming bond formation, which requires a precise substrate positioning, as in other cases^42^ (Supplementary Fig. 1). The three mutated residues are located next to each other (distances of C_α_ is ∼6 Å) lining the large binding pocket, but relatively far away (∼15-20 Å) from haem-Fe=O, assuming the absence of dynamic effects (**Fig. 1b**). Complete deconvolution of variant III starting from parental F87A entails 3! = 6 theoretically possible upward pathways, which we constructed by generating the 6 intermediate mutants (**Fig. 1c**). The key question is how these residues determine selectivity and activity, and whether they interact epistatically.

**Fig. 1.**
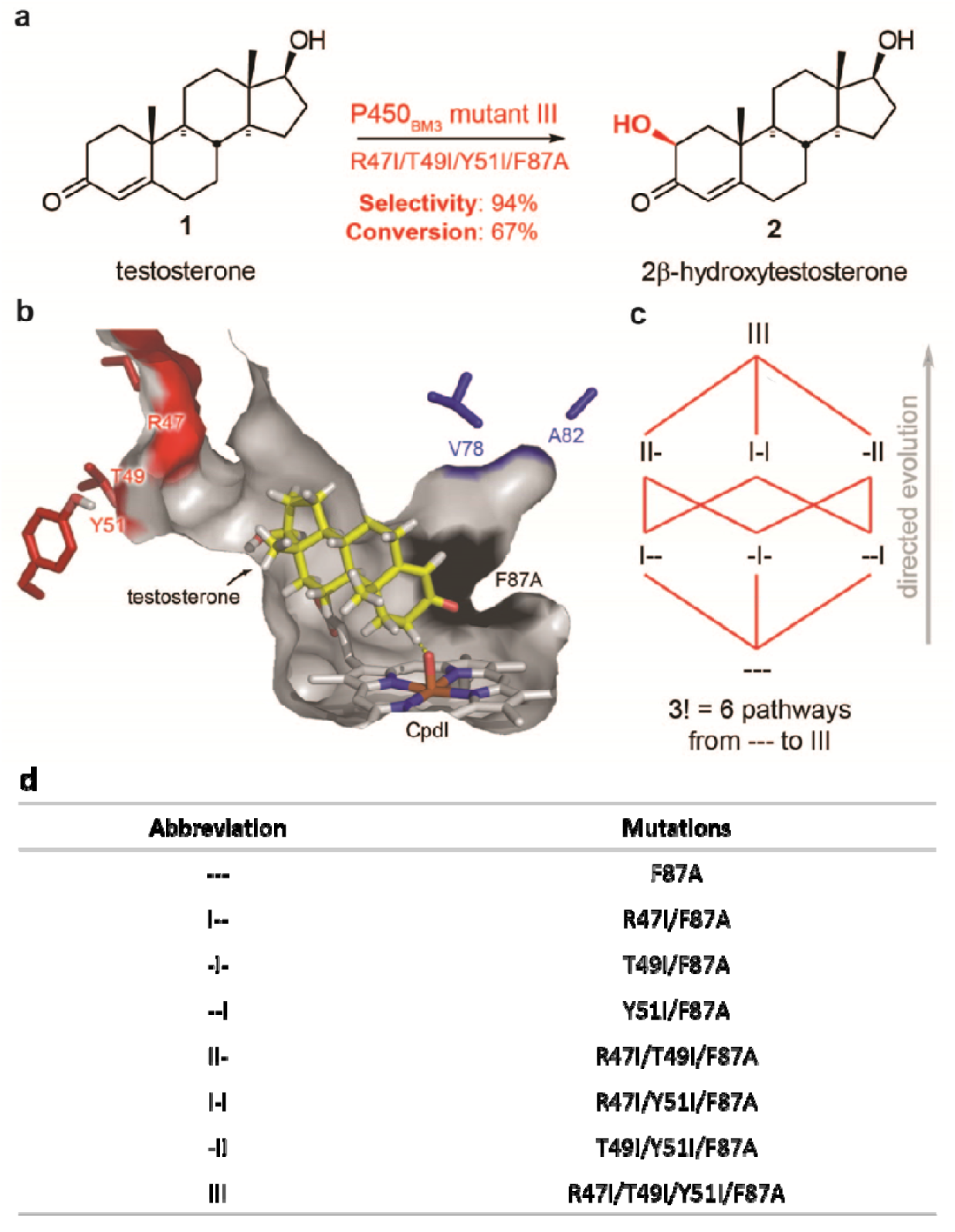
Model system based on P450_BM3_ as biocatalyst for the selective oxidation of a steroidal substrate. **a**) Testosterone (**1**) is selectively hydroxylated at position 2β (**2**) by mutant III (R47I/T49I/Y51I/F87A). An interactive figure of parental variant --- docked with **1** was created with Michelanglo^43^ (https://michelanglo.sgc.ox.ac.uk/r/p450) highlighting the mutated residues and secondary structures discussed in this work. The distances between the α-C atoms of the following pairs of residues are (Å): R47-T49 (7.0), R47-Y51 (13.4), T49-Y51 (7.0). Image and atom distances calculations obtained with PyMol Molecular Graphics System, V 1.5.0.4 (Schrödinger, LLC). **c**) The 6 possible evolutionary trajectories between parental mutant F87A (---) and “triple” mutant III involve three “single” mutants I-- (R47I/F87A), -I- (T49I/F87A) and --I (Y51I/F87A) as well as three “double” mutants II- (R47I/T49I/F87A), I-I (R47IY51I/F87A) and -II (T49I/Y51I/F87A). **d**) Mutant abbreviations.

All intermediate mutants were generated, overexpressed in *Escherichia coli* BL21-Gold(DE3) and purified (Supplementary Fig. 2). Parent F87A (---) and variant III were also included, resulting in a total of 8 enzymes. Using defined substrate and NADPH concentrations, multiple parameters were determined (Supplementary Note 1 and Table 1).

**Table 1:** Epistatic analysis of all possible mutational combinations among the III-derived mutants in pure form. This shortened data set originates from Supplementary Tables 2-8.

| Type<sup>a</sup> | Combination of mutations<sup>b</sup> | Resulting mutant<sup>b</sup> | Parameter<sup>c,d</sup> |  |  |  |  |  |  |
| --- | --- | --- | --- | --- | --- | --- | --- | --- | --- |
|  |  |  | Conv. | Sel. | TTN | TTF | PFR | NCR | CE |
| B | I-- + -I- | II- | A | +ME | -ME | -ME | -ME | -ME | +RSE |
| B | -I- + --I | -II | +SE | +SE | +ME | +ME | +ME | +ME | +SE |
| B | I-- + --I | I-I | +RSE | +SE | -ME | -ME | -ME | -ME | -SE |
| B | I-- + -II | III | +SE | +SE | A | +ME | +ME | +ME | +SE |
| B | II- + --I | III | +RSE | +SE | +ME | +ME | +ME | +ME | +ME |
| B | I-I + -I- | III | +SE | +SE | +ME | +ME | +ME | +ME | +SE |
| T | I-- + -I- + --I | III | +SE | +SE | +ME | +ME | +ME | +ME | +SE |

The most 2β-selective variants (∼67-91%) contain mutation Y51I (--I, I-I, -II and III), while the remaining ones proved to be 15β-selective, with substrate conversion being highest (35%) in mutants III and -II, and poor (∼6-10%) in the remaining ones (Fig. 2a). NADPH leaks without substrate in all mutants except -II and III, which display a respective 2- and 3-fold increased NADPH consumption upon **1** addition (Fig. 2b). Mutants -II and III also showed a respective ∼5- and ∼10-fold improvement in product formation rates (PFR) compared to the remaining variants (Fig. 2c), suggesting that variants -II and III have good coupling efficiency (CE). CE describes how well the reductase domain delivers electrons from NADPH via the flavin cofactors to the substrate in the haem domain. A low CE value indicates futile NADPH usage, resulting in the formation of reactive species during the catalytic cycle that can inactivate the enzyme^44^. Low CE values of 15-30% were found for all mutants, except for -II and III that display higher values of 37% (Fig. 2d). The total turnover number (TTN) is highest in mutants -II and III (Fig. 2e), whereas the total turnover frequency (TTF), PFR and NADPH consumption rate are highest in III (Fig. 2f).

**Fig. 2.**
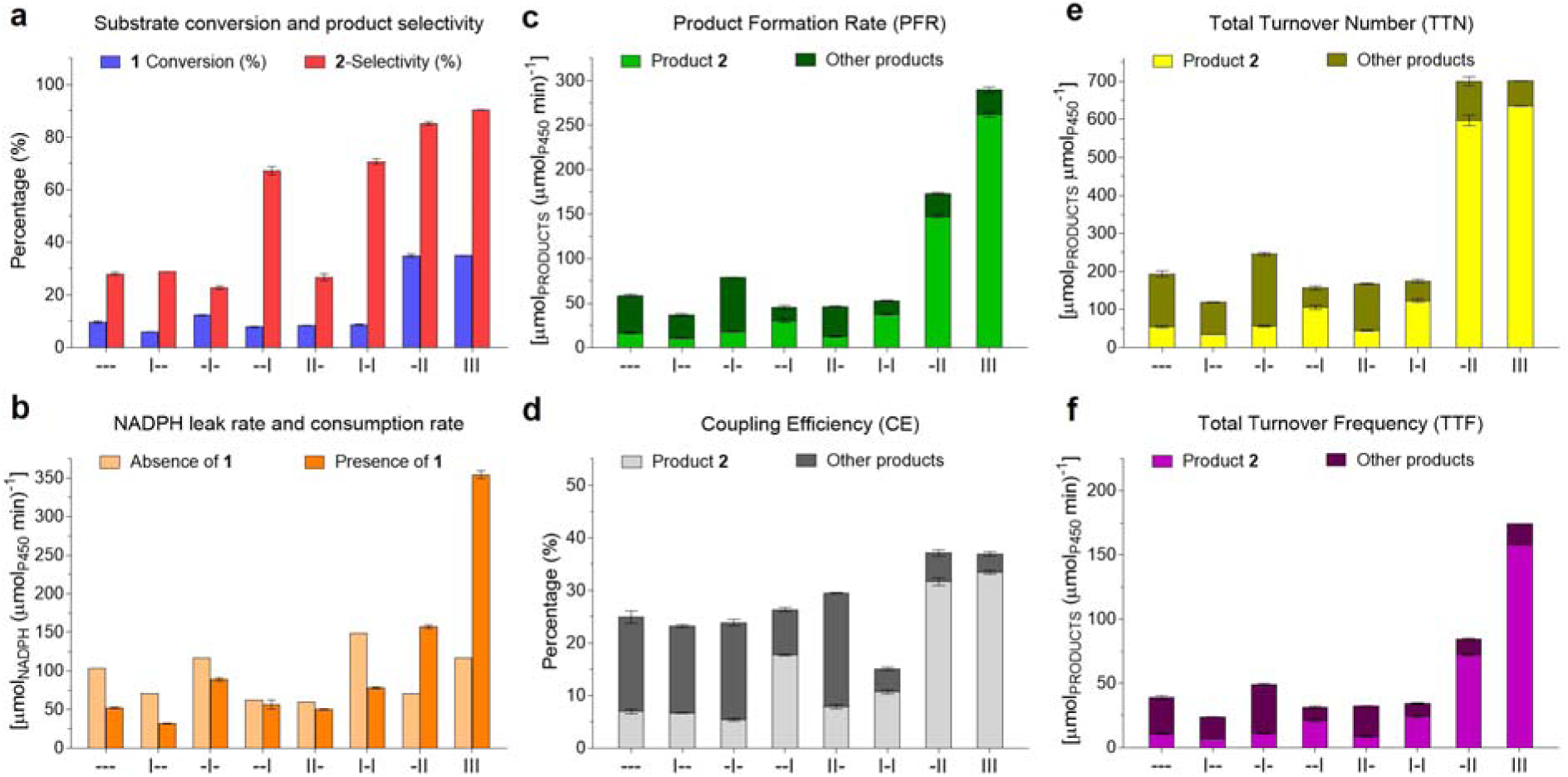
Multiple enzymatic parameters of deconvolution mutants. Selectivity and conversion (a) as well as coupling efficiency (d) data are reported in percentage. Initial rates of NADPH consumption (b) and PFR (c) were performed during the first 20 s upon addition of testosterone (**1**) substrate. Upon NADPH depletion, the reaction was stopped, and the samples analysed by HPLC. TTN (e) describes the total moles of products per mole of enzyme, and the TTF (f) normalises TTN by time after NADPH depletion. Other products mainly include mainly the 15β-alcohol and other regioisomers. The data represent the average ± s.e.m. of two independent experiments (*n*=2). See Fig. 1d for mutant abbreviations.

### Distal mutations enable conformational changes at the active site required for regioselectivity

To gain insights about the origin of selectivity and activity, we performed computational studies on all mutants. Given the identification of comparable reaction barriers for hydrogen-atom abstraction from C2 and C15 by using Density Functional Theory (DFT) calculations on truncated models (difference of <1.0 kcal per mol, see Supplementary Note 2 and Supplementary Fig. 3), we carried out MD simulations to analyse whether the binding pose of **1** in the active site determines the experimentally observed selectivity (Fig. 3 and Supplementary Note 3). Starting from parent ---, pose 2 (presenting C2 close to the catalytic Cpd I) and pose 15 (C15 close to Cpd I) generated from manual dockings are possible (Supplementary Figs. 4 and 5). Further analysis of these binding poses along MD simulations in --- indicate that substrate **1** in pose 15 explores near attack conformations (NACs)^45^ closer to the QM-predicted ideal transition state (TS) geometry for H-abstraction than in pose 2 (Fig. 3a), thus making pose 15 more productive towards 15β-hydroxylation. Introducing mutations R47I and/or T49I does not have any effect on selectivity (Fig. 3b-c, e), i.e., the selectivity is retained due to the catalytically competent conformation inherent in pose 15 along MD simulations (pose 2 adopts a reduced number of catalytically competent conformations). However, the picture completely changes when mutation Y51I is introduced: the substrate bound in pose 15 becomes unstable and leaves the active site in 1 out of 3 replicas (ca. >15 Å C2··O distances explored, Fig. 3d), whereas pose 2 is highly stabilized and explores short C2··O distances for the incipient C-H eventually leading to 2β-hydroxytestosterone in 2 out of 3 replicas (Fig. 3d and Supplementary Figs. 6 and 7). As experimentally determined, 2β-selectivity is retained in variants I-I, -II and III that contain mutation Y51I (Fig. 3f-h). This is even more dramatic in variant III, in which pose 15 is highly unstable and **1** rapidly rotates to position C2 close to the catalytic Cpd I for 2β-hydroxylation (Supplementary Video 1).

**Fig. 3.**
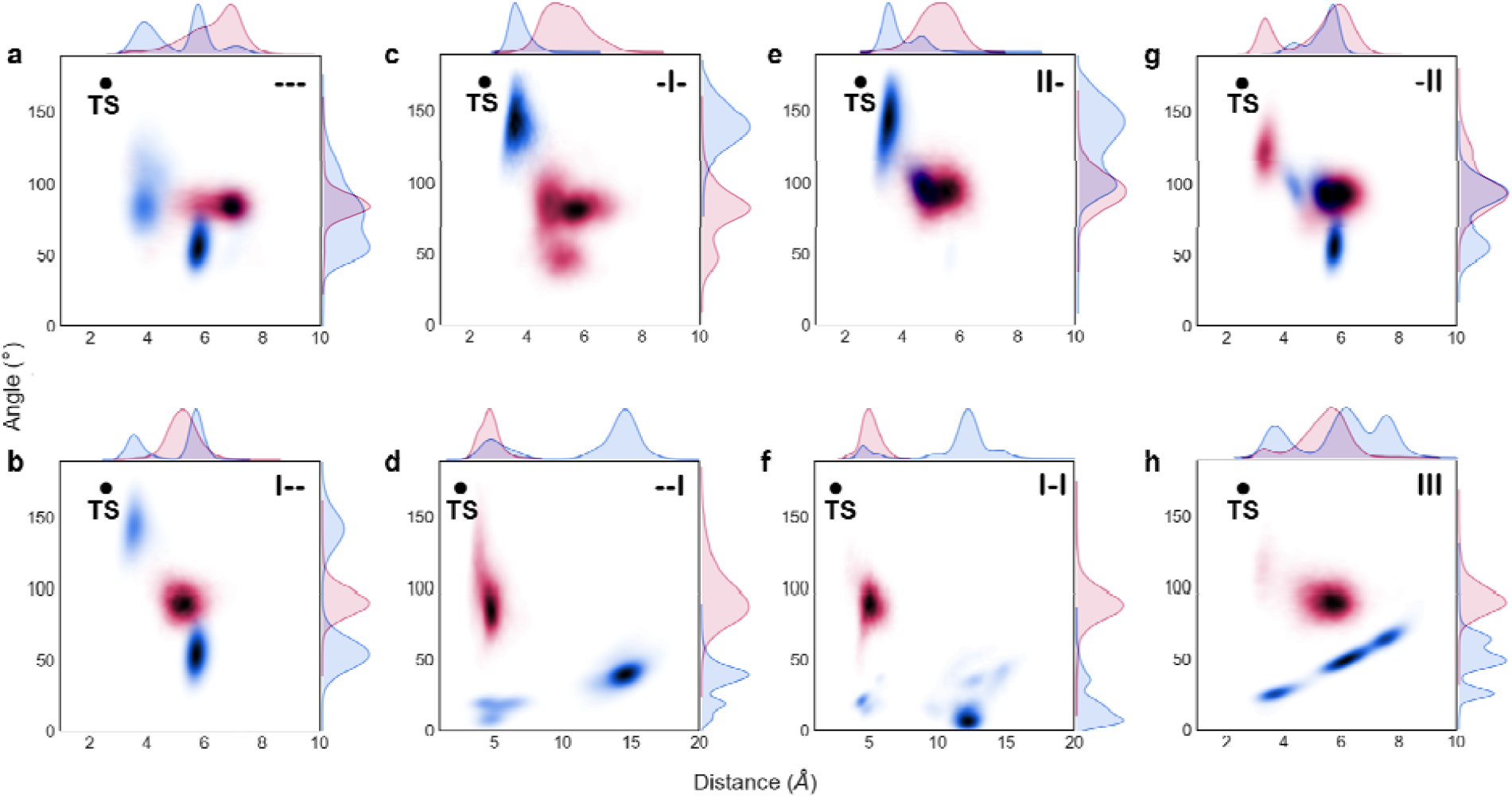
Conformational population analysis of key geometric parameters for hydroxylation. Distances determined between the oxygen atom of haem-Fe=O and the C-atom (C2 or C15) of **1** (x-axis) and angles formed by O(Fe=O) – (**1**)-H(C2/15) – (**1**)- C(2/15) (y axis) from the first replica of the MD dataset of all mutants (see Supplementary Figs. 6 and 7 for additional replicas). Geometric parameters measured for C-2 and C-15 are shown in red and red blue, respectively. The ideal distance and angle for the TS (black dot) corresponds to the Density Functional Theory (DFT) optimized geometry for the C–H abstraction by haem-Fe=O using a truncated computational model (Supplementary Note 2). TS: Transition State. See Fig. 1d for mutant abbreviations.

Notably, substrate rotation inside the active site pocket is only observed in mutant III, which presents a substantially wider active site pocket as compared to the other variants: the active site volume in the --- variant is 89 Å^3^, which is expanded to 235 Å^3^ in III (Supplementary Fig. 8). We hypothesised that in all variants, except III, selectivity must be determined by the orientation adopted by the substrate while accessing the haem cavity. Recently, Mondal et al. characterized the substrate recognition and binding pathway in related P450cam using MD simulations, showing the formation of a single key channel in which the substrate needs to reside in a long-lived intermediate state before reaching the catalytic iron-oxo species^46^.

To reconstruct the substrate binding process in P450_BM3_, we placed substrate **1** in the bulk solvent and started unbiased MD simulations followed by accelerated Molecular Dynamics (aMD) simulations (Supplementary Note 4). Among independent MD and aMD trajectories for all variants (250 ns MD + 750 ns aMD), only a single trajectory by mutant I-- was observed to be productive where the substrate reached the haem active site (Fig. 4b). We observed a two-step binding mechanism in this trajectory: first, the carbonyl moiety of **1** enters channel 2a (Fig. 4a) and stays between β1-4 strand, where residues R47I, T49I, Y51I are located, forming a long-lived substrate-enzyme bound intermediate. There, substrate **1** can reorient, although its access to the active site is restricted by the β4 sheet that acts as a gate. Second, a network of coupled conformational changes occur simultaneously: G helix adopts a bend conformation, which impacts F helix and β1 sheet conformation, and in turn shifts B’ helix and retreats β4 sheet, allowing **1** progression towards the catalytic centre (Supplementary Video 2). This 2-step mechanism is similar to what Mondal et al. observed in P450cam^46^.

**Fig. 4.**
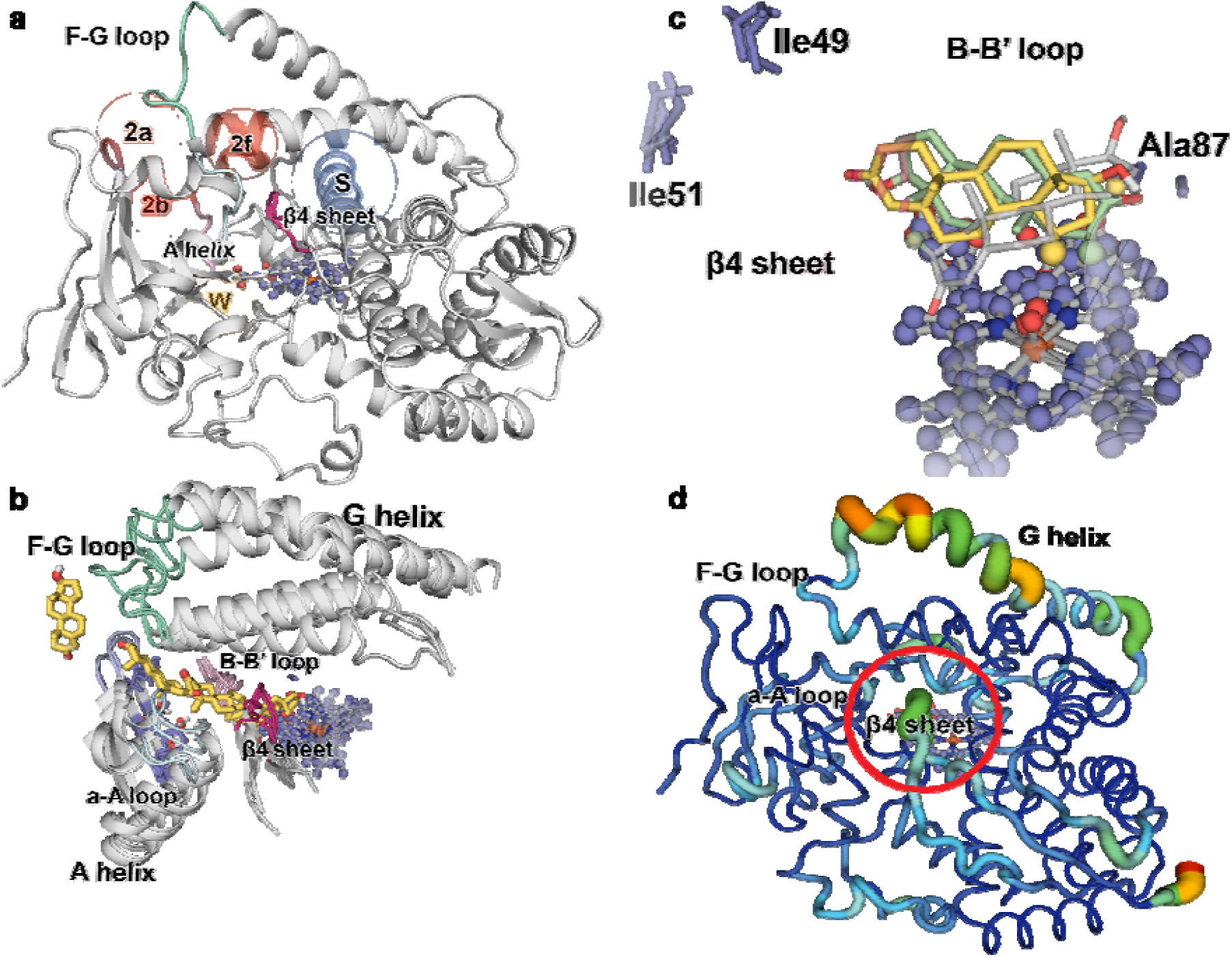
Secondary structural elements determining regioselectivity and activity. a) Spheres indicate the channels observed in the WT crystal structure (PDB: 1FAG) and in the mutants of our MD simulations (red and blue colour). b) Trajectories of **1** towards the active site of mutant I-- and binding of **1** above the haem. c) Rotation of **1** from pose15 to pose2 in the active site of mutant III. β hydrogens belonging to C2 and C15 atoms are depicted in pale green and yellow colour, respectively. d) Principal Component Analysis (pc2) of mutant III (APO). The thickness of the line is proportional to the motion and the colour scale varies from blue (minimum motion) to red (maximum motion). The β4 sheet is highlighted with a red circle. Standard nomenclature for channels^48^ and secondary structure elements^49^ is used.

Our aMD simulations show that the orientation of the substrate when accessing the catalytic site during the second step of the binding pathway dictates selectivity. In the productive trajectory corresponding to mutant I--, **1** accesses the haem with the correct orientation for 15β-hydroxylation (Fig. 4b and Supplementary Fig. 9). In this case, residue Y51 establishes a hydrogen bond with the carbonyl group of **1** (Supplementary Fig. 10), constraining the substrate in a such way that it can only progress into the active site pocket pointing its C15 ahead towards haem-Fe=O^47^. Thus, Y51 is instrumental in promoting the observed C15-selectivity in ---, and in I--, -I- and II-variants. Additionally, the higher C2-selectivity observed in variant III occurs due to the flipping and motion of the β4 sheet, destabilizing pose 15 while favouring pose 2 (Fig. 4c).

Interestingly, the analysis of the most relevant conformational changes in each independent variant through Principal Component Analysis (PCA) indicates that the most active mutant III shows the highest flexibility of the β4 sheet (Fig. 4d). These flexible regions, responsible of controlling substrate binding as described above, influence activity, as mutant III shows the highest TTF, NADPH consumption rate and PFR numbers. Thus, favouring a more efficient substrate binding in a catalytically competent pose increases enzyme TTF, while NADPH leak is reduced due to a more efficient interaction between the substrate and the catalytically active Fe=O species once generated.

### Pervasive epistatic effects on multiple parameters are cooperative

Based on Tokuriki^11^ and Bendixsen et al.^50^, non-additive mutational effects can occur in different forms (Fig. 5), which can be calculated with additivity equations (Supplementary Note 5). Aiming at exploring the existence of epistatic effects in an effective manner, we developed a Python-based computational program to automatically determine the type and intensity of amino acid interactions among all possible mutational combinations (Supplementary Note 6).

**Fig. 5.**
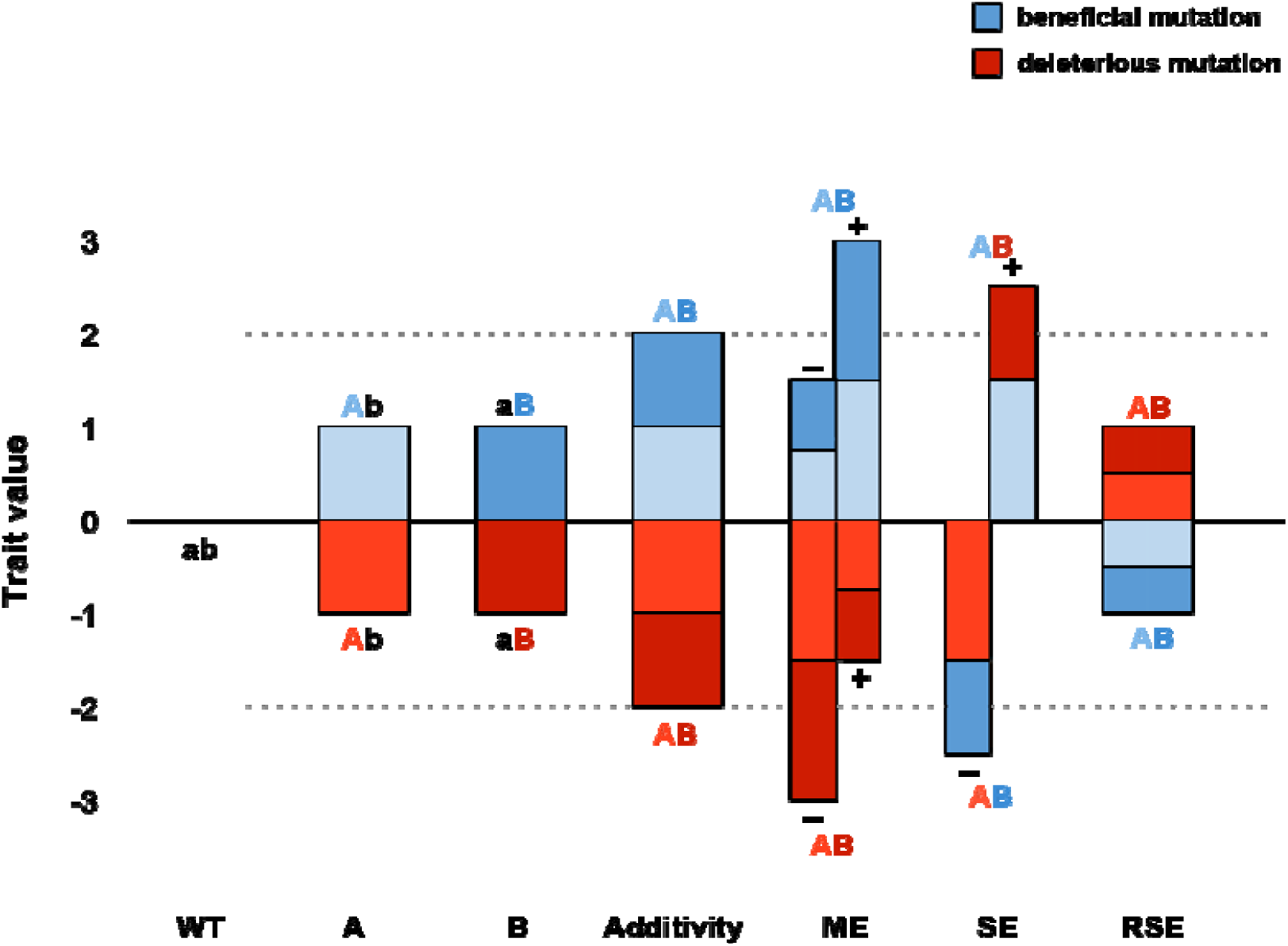
Non-additive effects in single mutations A and B. Epistatic effects emerge when the result of combining two individual mutations (or sets of mutations) is non-additive (i.e., the fitness value after combining mutations A with B to generate a doubly mutated AB variant is not equal to the sum of the individual A and B contributions). Epistatic effects can be positive/synergistic or negative/antagonistic depending on the fitness trait. They occur in the form of: (i) magnitude epistasis (ME) if both the single mutations A and B are beneficial for a fitness trait and they produce a greater-than-additive fitness improvement when combined together in mutant AB; (ii) sign epistasis (SE) if one mutation A is deleterious on its own but can enhance the beneficial effect of another mutation B if combined into AB; (iii) reciprocal sign epistasis (RSE) if both mutations A and B are deleterious alone, but they produce a beneficial effect when combined in AB (or if both are beneficial if they produce a deleterious effect).

We quantified all amino acid interactions among all 6 trajectories leading from parent --- to mutant III for multiple parameters (Table 1). Most potential combinations on substrate conversion show synergistic effects, with 4 (57 %) and 2 (29 %) cases of sign epistasis (SE) and reciprocal sign epistasis (RSE), respectively, and one single case of additivity (Supplementary Table 2). For 2β-selectivity, all interactions are synergistic, with most of them showing positive sign epistasis and only one case of positive magnitude epistasis (combination of R47I and T49I). For example, the combination of the single mutations R47I, T49I, and Y51I (in parent mutant ---) is expected to contribute -3.45 ± 0.25 kJ per mol. The two former mutations confer 15β-selectivity in mutants I-- and -I-, while the latter one induces 2β-selectivity in variant --I. Yet the experimental value of the 15β-selective mutant III yields 5.6 ± 0.0 kJ per mol, which represents a difference of about 9 kJ per mol between the experimental and theoretical values (Supplementary Table 3). TTF, PFR and NADPH consumption rate show essentially the same type of epistatic effects, i.e., all interactions generally yield magnitude epistasis with 5 (∼70 %) and 2 (∼30 %) being positive (+ME) and negative (-ME), respectively (Supplementary Tables 4-6). Very similar epistatic effects are seen for TTN in all mutational combinations, except for one case of additivity (Supplementary Table 7). Finally, coupling efficiency shows 6 cases of synergistic epistatic effects and 1 case of antagonistic epistatic effects (Supplementary Table 8). These results indicate that an efficient formation of 2β-hydroxytestosterone requires pervasive cooperative effects among R47, T49 and R51 regardless of mutational combination.

### Conformational dynamics shape the evolution of fitness landscapes

The complete deconvolution of a multi-mutational variant enables the exploration of all possible pathways from parental enzyme to the evolved mutant, thus determining a full multidimensional fitness landscape. To explore the step-wise accessibility in the evolution of parental --- towards III, we constructed a fitness “pathway” landscape^51^, which for the first time focuses on activity *and* on selectivity (Fig. 6a). This system is a 4-dimensional surface (3 sets of mutations as independent vectors and ΔΔG^‡^ as the dependent variable obtained from the experimental selectivities). Two kinds of trajectories can be noted: those lacking local minima (favoured) and those characterized by at least one local minimum (disfavoured). Pathways 1-4 are characterized by a decrease in both selectivity and activity at the first step, indicating that they are evolutionarily disfavoured (pathway 3 is highlighted in red). Pathway 5 (highlighted in green) and 6 are favoured because --I enables conformational changes in the active site and has implications on the substrate binding (as discussed above). In the two latter pathways, activity improves slightly in the evolution of --- towards --I (TTF = 11 → 21) at the first step, but at the second and third steps of pathway 6 it increases significantly towards -II and III (TTF = 21 → 72 → 158). This is due to the β4 sheet that shows an increased flexibility in the most active mutants -II and III, highlighting the key role of β4 sheet for activity (Fig. 6b). Interestingly, when NADPH consumption rate is additionally considered, pathways 3 and 4 become accessible (Supplementary Note 7 and Supplementary Fig. 11); however, the improvements are not as pronounced as in activity and selectivity.

**Fig. 6.**
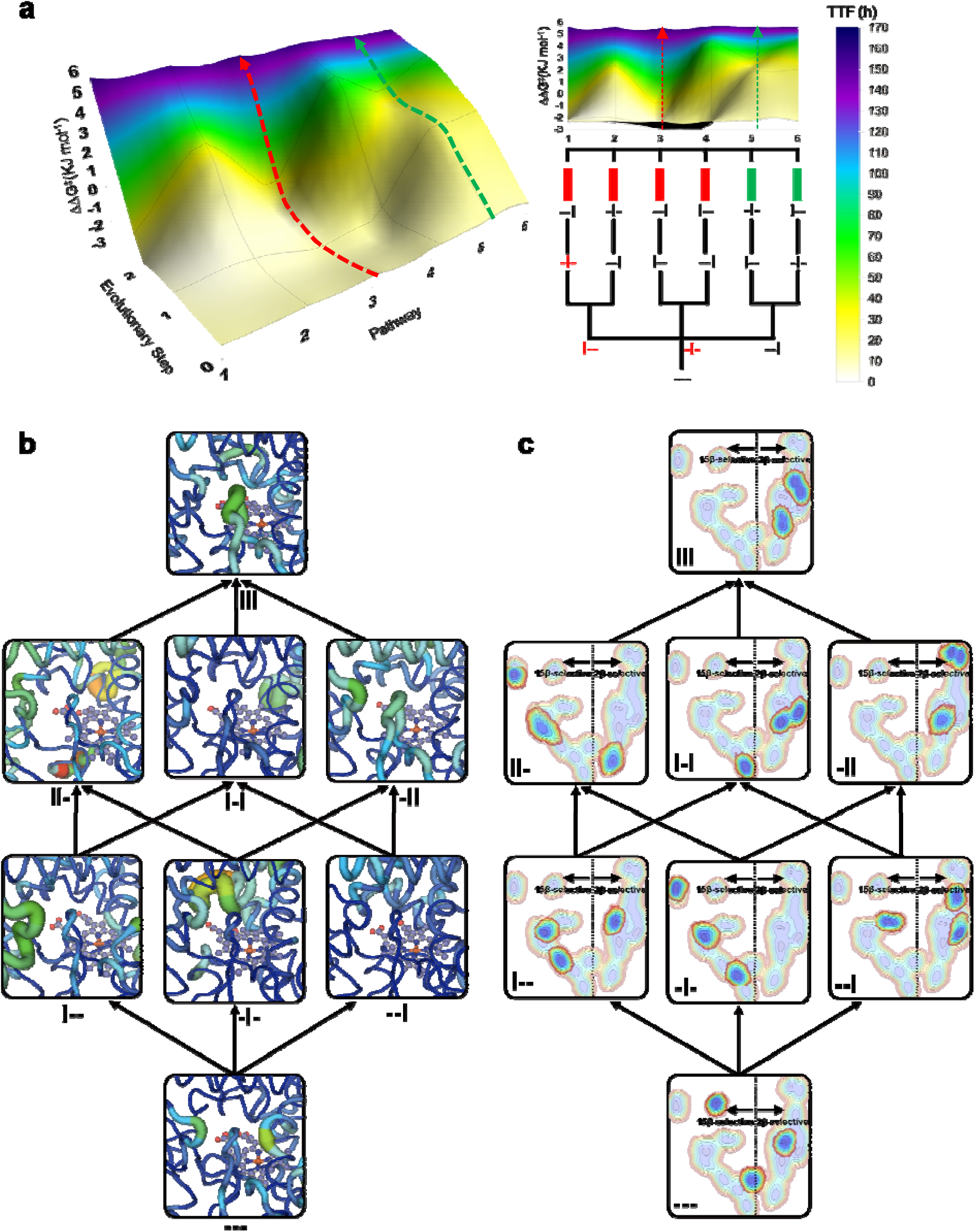
Stepwise evolution of multiple functions and conformational dynamics. a) Multiparametric fitness pathway landscapes of the 6 evolutionary pathways leading from parent mutant --- upward to mutant III are shown in 3D (left) and frontal (right) format (green and red arrows indicate examples of favourable and unfavourable pathways, respectively). Fitness is defined by activity as total turnover frequency (TTF) displayed as heat-maps from 0 (white) to 170 (blue) as well as 2β-selectivity as ΔΔG^‡^ in kJ per mol (y-axis) at each evolutionary step (z-axis) for each pathway (x-axis). Green and red bars indicate favoured and disfavoured pathways, respectively. The mutant in red represents steps with disfavoured energy, i.e., the point where the pathway is blocked. Source data are listed in Supplementary Table 1 and pathway analysis in Supplementary Tables 9 and 10. b) Progression of the β4 sheet flexibility along the 6 pathways as revealed by PC analysis (pc2) of the substrate-free simulations of mutated enzymes analysed separately (see complete haem domains in Supplementary Fig. 12). The thickness of the line is proportional to the motion and the colour scale varies from blue (minimum motion) to red (maximum motion). c) Evolution of the conformational dynamics along the 6 pathways and its connection to 2β- or 15β-selectivity. The analysis of the global conformational dynamics of the substrate-free simulations of mutated enzymes, as shown by pc1/pc3, indicate that 2β- and 15β-selective mutants explore conformations lying at positive and negative values of pc1, respectively. Color scale varies from red (less populated) to blue (more populated). See Fig. 1d for mutant abbreviations.

To identify the most important conformational changes in all evolutionary pathways, and to describe how distal mutations influence them, we performed extensive MD simulations in the absence of substrate of each variant and applied the dimensionality reduction technique PCA^28^ to the whole dataset (Fig. 6c and 7a). A conformational population analysis resulting from all the accumulated simulation data was generated in terms of principal components (PC) 1 and 3, which describe the first and third most important conformational differences among all variants (for PC2 see Supplementary Fig. 13). Notably, a clear distinction between 2- and 15β-selective mutants is revealed through their separation with respect to PC1 (x-axis), suggesting that changes in selectivity are linked to the impact that the introduced mutations have on the enzyme conformational dynamics (Fig. 6c). These conformational changes mainly involve the G helix, the F-G loop, the β1 hairpin and the B’ helix (located at the entrance of the 2a/b channels) as well as the a-A loop and the β4 sheet (located at the entrance of the 2f channel) (Fig. 7b). In variant -I-, the channel 2a has a narrower substrate access entrance due to a closed state of the F-G loop (*ca.* 9.3 Å determined between the Cα of R47 and N192). Conversely, the combination of mutations introduced in III favours an open conformational state of the same F-G loop (*ca.* 12.4 Å measured between the Cα of I47 and N192), enlarging the access channel 2a, which is mainly responsible for allowing access to the enzyme binding pocket (Fig. 7c). Indeed, the area surrounding the access channel 2a in mutant III is calculated to have a volume of 140 Å^3^ with respect to 44 Å^3^ in -I-.

**Fig. 7.**
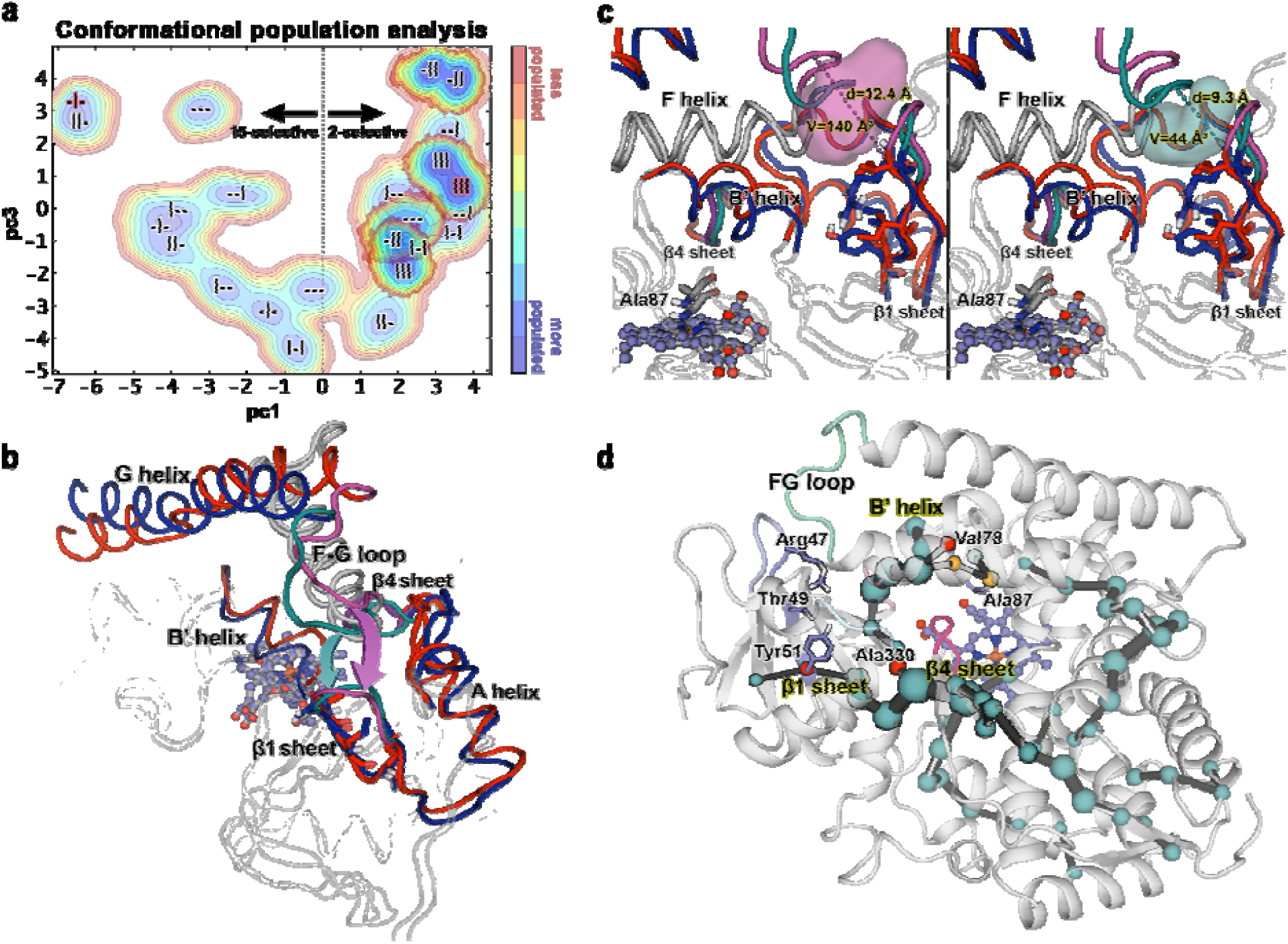
Analysis of the conformational dynamics of deconvolution mutants. a) Conformational population analysis built from the combined PCA of all substrate-free simulations of mutants. The conformational populations of 2β-selective mutants -II and III are highlighted. All replicas (3/3) of III and -II, 2/3 replica of I-I and I-- lie on positive values of pc1 (10 out of 12 replicas of 2β-selective mutants). All replicas (3/3) of -I-, 2/3 replicas of ---, I-- and II-show negative values of pc1 (9 out of 12 replicas of all 15β-selective mutants). b) Overlay of the conformational changes involved in PC1. c) Zoom of (b) showing the channel 2a cavity of mutant III (left panel) and -I- (right panel). In (b) and (c) Mutant -I- is shown in blue, whereas the evolved mutant III is coloured in red. The F-G loop, β1 sheet and channel 2a cavity are highlighted in teal and magenta for -I- and III, respectively. d) Analysis of the most important correlated motions of mutant --- by means of shortest path map (SPM). Mutational hotspot used in this or precedent work^41^ that appear in the SPM are highlighted with red spheres, whereas positions adjacent to mutational hotspot are highlighted with orange spheres. See Fig. 1d for mutant abbreviations.

To further study the link between epistasis and conformational dynamics, we applied the Shortest Path Map (SPM) analysis^21^ (Supplementary Note 9) using the accumulated 1.8 μs MD simulation performed on parent --- in the absence of substrate. SPM considers the different conformations that the enzyme samples along the MD simulation, and identifies which residues are those that are more important for the observed conformational changes, which in this case are associated with different selectivities and activities^21^. In the parent --- enzyme, the generated SPM identifies residues Y51 as well as V78 and A330, known from earlier studies^38^, to be important for enzyme activity and to be interconnected in terms of Cα correlated movements, thus highly contributing to the enzyme inactive-to-active conformational interconversion. This highlights why these three distal positions are found to be key during the evolutionary pathway for improving catalysis, in line with what we observed for the laboratory-evolved retro-aldolases^21^. Importantly, the SPM also describes strong connections between all the five-stranded β1 sheet with the β4-2 strand and the B’ helix, which we showed to be crucial for substrate binding and gating (Fig. 7d). This long-distance communicating pathway between β1 and β4 sheets directly relates the mutated positions on β1-4 strand (positions R47, T49 and Y51) and the increased flexibility of the β4 sheet. This shows how evolutionary pathways take advantage of networks of residue-residue interactions to fine tune the conformational dynamics along the evolutionary pathways for improving enzyme function.

## Discussion

The identification of epistasis and its molecular mechanism are crucial for understanding protein function, but these are hardly ever explored in directed or natural evolution studies of multiparametric optimization. To the best of our knowledge, only the study by Hartl and colleagues on fitness landscapes of drug resistance towards trimethoprim fits such a description, in which activity, binding and folding stability were shown to depend to some extent on epistasis^52^. Other studies dealing with directed evolution of activity and selectivity in plant sesquiterpene synthases^53^ and P450_BM3_^54^ do not consider epistatic effects, while “stability-mediated epistatic effects” were observed in a P450_BM3_ study^55^ some years later^5^.

The construction of fitness landscapes using a single catalytic parameter has been reported in two main research areas. While such landscapes have revealed that usually many pathways are accessible in laboratory evolution of enzymes as catalysts in organic chemistry^4,7,51,56^, different conclusions have been made in evolutionary biology^8–11^. In the present study, we observed that there are no direct accessible pathways to activity and that only a few trajectories (2/6) are accessible to both selectivity and activity. Interestingly, the two accessible pathways for selectivity correspond to mutation Y51I. The addition of mutation R47I has almost no effect on activity and selectivity, while mutation T49I, which is closer to Y51I, significantly improves both parameters as it alters the enzyme conformational dynamics. T49I and Y51I enhance the flexibility of the β4 sheet, and both combined with R47I reshape the active site for enhanced 2β-hydroxylation. The triple mutant III excels in all parameters compared to all double and single mutants. Unexpectedly, upon going from the “parent” enzyme --- to mutant III, cooperative effects at each step in the evolution of activity and selectivity remain pervasive. Residues R47I, T49I and Y51I are located at the entrance of a long substrate channel far away from the active site^41^. Since the mutated residues were not observed to interact directly with the substrate in our MD simulations, we propose that the observed epistatic effects, which are mediated by long-range interactions, can occur via one main mechanism: *Direct effects between mutations, but no direct interaction between the substrate and the mutations*^11^.

Our computational exploration of the mutation-induced conformational changes on F87A variants provide key insights concerning the importance for P450_BM3_ evolution towards more active and selective variants. These simulations have shown, for the first time, that *activity is dictated by the flexibility of the* β*4 sheet*, which acts as a gate and modulates substrate access to the catalytic haem pocket for efficient hydroxylation. Our simulations also highlight the key role of the F-G loop in open-close conformational transitions involved in substrate binding, as shown for P450_BM3_ by Shaik^57^ and P450_PikC_ by Houk and Sherman^58^. The substrate binding simulations show that *selectivity is dictated by how the substrate is oriented when accessing the haem pocket* through the β4 sheet. By tuning the open/close conformational states of the F-G loop and the β1 hairpin, the substrate access channels are altered, which impact substrate orientation and thus selectivity. This rich conformational heterogeneity observed for P450_BM3_ important for substrate binding is in line with previous reports^57^ and also with the selective stabilization of discrete conformational states of P450_CYP119_ and P450_PikC_ upon ligand binding^58,59^. However, our simulations contrast to what was previously observed in P450_cam_, which does not depend on open/closed conformational changes of the F, G helices and loop for allowing substrate binding^46^. It should be also mentioned that P450_cam_ complexed with its redox partner adopts an open conformation that stabilizes the active site key for the proton relay network^60^.

Using SPM analysis, the most important positions that participate in the open/closed conformational conversions that dictate selectivity and activity were identified. Of relevance is that the key residue Y51 found to be essential for both activity and selectivity in this study, is contained in the SPM path, as well as the previously described V78 and A330 positions^38^. SPM also highlights a long-distance communicating pathway between β4 and β1 where positions R47, T49, and Y51 are located, which is exploited along the evolutionary pathway for altering BM3 protein function.

This study evidences that epistasis is intrinsically linked to conformational dynamics, which fine-tunes multiple functions in a protein involved in secondary metabolism. Our findings on the conformational changes connected to CYP activity and selectivity and residue networks that modulate such conformational conversions can be expected to facilitate future rational evolution of these enzymes.

## Acknowledgments

Support from the Max-Planck-Society and the LOEWE Research cluster SynChemBio is gratefully acknowledged. This study was also supported in part by the European Research Council Horizon 2020 research and innovation program (ERC-2015-StG-679001, S.O.), Spanish MINECO (project PGC2018-102192-B-I00, S.O.; and Juan de la Cierva - Incorporación fellowship IJCI-2017-33411, M.G.B.), UdG (predoctoral fellowship IFUdG2016, L.D.), and Generalitat de Catalunya AGAUR (SGR-1707, S.O.; and Beatriu de Pinós H2020 MSCA-Cofund 2018-BP-00204, M.G.-B.).

## Author contributions

C.G.A.R., S.O. and M.T.R. conceived the project; C.G.A.R., A.L. and S.H. created and purified mutants and measured their activity; P.L. and M.F. wrote the Python code for automated calculation of epistatic effects; J.S. constructed and analyzed the fitness pathway landscapes; L.D. performed the computational modeling and analysis with support and guidance from M.G.B. and S.O.; all authors revised and approved the manuscript.

## Additional information

Supplementary information is available in the online version of the paper.

## Methods

### Chemicals, materials and software

All commercial chemicals were purchased with the highest purity grade (e.g., HPLC) from Sigma-Aldrich (St. Louis, US) unless otherwise indicated. For protein purification, lysozyme and DNase I was purchased from Applichem (Darmstadt, Germany). For PCRs, KOD Hot-Start DNA Polymerase was obtained from Novagen (Merck, Darmstadt, Germany). Restriction enzyme *DpnI* was bought from New England Biolabs (Ipswich, US). The *E. coli* BL21-Gold(DE3) strain, obtained from Novagen (Merck-Millipore) and generally cultured in lysogeny broth (LB) with 50 μg/mL kanamycin (^Kan50^) as marker (LB^Kan50^), both obtained from Carl Roth, was used for transformation of site-directed and saturation mutagenesis reactions as well as for protein over-expression experiments. According to standard molecular biology protocols, electro-competent *E. coli* cells were prepared using 10% glycerol (Applichem) and transformed with the corresponding plasmids using a “MicroPulser” electroporator (BioRad, Hercules, US) following the manufacturer’s instructions. For site-directed and saturation mutagenesis experiments, oligonucleotides were purchased from Metabion (Martinsried, Germany) and Integrated DNA Technologies (Iowa, US). Plasmid isolation kit (mini-preparation) was ordered from Zymo Research (Irvine, US) and QIAGEN (Hildesheim, Germany). DNA sequencing was conducted by GATC Biotech (Konstanz, Germany). The analysis of sequencing reads was performed using the commercial software MegAlign from DNASTAR Lasergene version 11 (Madison, US) and the freeware ApE plasmid editor version 2.0.44 by Wayne Davis. The software used for constructing the fitness pathway landscapes is Surfer version 8 (Golden, US) and for the thermodynamic cycles is GraphPad Prism version 6 (La Jolla, US).

### P450_BM3_-based oxidation reactions using purified enzymes

Biotransformation reactions were performed as described earlier^36^. Briefly, reactions were performed in 2.2 mL MTP format by resuspending the thawed cells in 600 μL of reaction mixture, followed by addition of 6 μL testosterone [stock: 100 mM (DMF); final conc. 1 mM (1%)] and incubation for 24 h at 25°C, 220 rpm using gas permeable seals in the same orbital shaker described above. The reaction mixture consisted of 100 mM KPi buffer pH 8.0, 100 mM glucose (Applichem), 10% glycerol (Applichem), 1 mM NADP^+^ (Merck-Millipore or Applichem), 1 U/mL glucose dehydrogenase (GDH-105) obtained from Codexis (Redwood City, US), 5 mM EDTA and 50 μg/mL kanamycin. The reaction was stopped by adding 350 μL of ethyl acetate using a Tecan robotic system (Männedorf, Switzerland) equipped with a liquid handling arm (LiHA), which was controlled using Gemini software V3.50, followed by centrifugation (10 min, 4,000 rpm, 4°C). The organic phase was extracted using the same robotic system but with the multi-pipette option (Te-MO), transferred to 500 μL MTPs (Nunc, Roskilde, Denmark) and left unsealed for evaporation in the fume hood overnight. The dried samples were resuspended in 150 μL acetonitrile and passed through a PTSF 96-well plate filter to remove solid particles (Pall, VWR, Germany) into a new 500 μL MTP (Nunc). The MTPs, which were closed using silicon lids for the corresponding plates, were stored at 4°C prior to screening.

### Steroid hydroxylation screening by HPLC

A LC-2010 HPLC system (Shimadzu, Japan) equipped with four MTP racks was used employing a reverse-phase “250 Eclipse XDB” C18 column of 250 mm (1.8 μM size particle) together with a corresponding pre-column bought from Agilent (Waldbronn, Germany) as stationary phase and installed in the oven at 40°C. The mobile phase was composed of a mixture of high-purity water generated from the local deionized water supply using a TKA MicroLab water purification system, acetonitrile (CH_3_CN) and methanol (MeOH). For testosterone (**1**), a program of 8 min based on a CH_3_CN:MeOH:H_2_O mixture was used: 0 →3 min (15:15:70), 3 →5 min (20:20:60), 5 → 6 min (30:30:40), 6 →7 min (15:15:70). This protocol allows the separation of >14 oxidation products of **1**. The retention times of the known and unknown compounds can be found elsewhere earlier^36^. Data analysis and plotting of screening results were performed using MS Excel 2016.

### Large-scale protein expression and purification

The P450_BM3_ mutants were inoculated into 4 mL LB^Kan50^ broth and cultured overnight at 37°C, 220 rpm. The overnight culture (4 mL) was transferred into 200 mL TB^Kan50^ in 500 mL shaking flasks. The cultivation continued at 37°C, 220 rpm for 2∼3 h until the OD_600_ reached ∼0.6-0.8, then IPTG was added to a final concentration of 100 μM and the temperature was reduced to 25°C. After 20 h expression, the cells were harvested by centrifugation at 4000 rpm, 4°C for 15 min. The cell pellets were stored at -80°C until further processing. The cell pellets were dissolved in buffer (50 mM KPi, 800 mM NaCl, pH 7.5) and disrupted by sonication under an ice bath. The collected lysate was centrifuged for 45 min at 11,000 rpm at 4°C and the obtained brownish-red supernatant was filtered to sterility with a 0.45 μm filter. The lysate obtained was loaded onto the pre-equilibrated nickel affinity column (HisTrap FF, 5 mL, GE Healthcare) with loading buffer (50 mM KPi, 800 mM NaCl, pH 7.5, 2 mM L-histidine). The column was first washed with 10 column volumes loading buffer, followed by gradient elution using an L-Histidine buffer (50 mM KPi, 80 mM L-histidine, pH 7.5) until complete protein elution. Columns were stripped and recharged between each mutant to avoid cross contamination. A flow rate of 5 mL/min was used and all fractions showing adsorption at 417 nm were collected. Proteins from the flow through were pooled and the buffer was exchanged to 25 mM KPi (pH 7.5) by ultrafiltration using a 50 kDa Amicon Ultra centrifugal filter (Merck-Millipore), and then concentrated to 5 mL. To remove the bound endogenous fatty acid, gravity-flow protein purification with Lipidex 1000 (Perkin Elmer) chromatography was conducted. 10 mL of Lipidex resin stored in methanol was used for column packing, which was subsequently washed with 10 column volumes of water and 10 column volumes of buffer (25 mM KPi, pH 7.5). After that, the protein was applied onto the column. The column was then capped to leave the protein in contact with resin at room temperature for 1 h, allowing hydrophobic compounds to bind to the resin. The protein was completely eluted from the Lipidex resin with buffer (25 mM KPi, pH 7.5) and the column was cleaned with at least 10 column volumes of methanol. The purified protein was pooled, and the buffer was exchanged to 100 mM KPi (pH 8.0) by ultrafiltration using a 50 kDa Amicon Ultra centrifugal filter, and then concentrated to 1 mL and stored in -80°C for further use. An aliquot was thawed at room temperature and enzyme concentration was determined by CO difference spectrum analysis prior to usage. The enzyme concentration determined for all intermediate mutants is shown in Supplementary Table 1.

### Determination of kinetic parameters using isolated enzymes

The kinetic experiments were performed using a JASCO V-650 spectrophotometer (JASCO International CO., LTD, Japan) equipped with a PAC-743 Peltier temperature control unit and UV-Vis-NIR Spectra Manager software II. All assays were performed in 100 mM potassium phosphate buffer (pH 8.0) at 25°C using quartz cuvettes adapted for magnetic stirring (900 rpm). The initial rate based on NADPH consumption was determined by measuring NADPH depletion monitored at 340 nm (ε = 6.22 mM-^1^ cm-^1^).

The NADPH concentration 0.24 mM and P450_BM3_ mutant concentrations (10∼100 nM) were employed in the reaction mixture. Reactions were started by adding testosterone from the stock solution in DMF with a final concentration of 0.2 mM. The concentration of DMF was 1% (v/v) for all measurements. Initial reaction rates were calculated from the first 20 s of the reaction. Due to uncoupling reactions, where NADPH is consumed without substrate hydroxylation, the initial rate calculation for different mutants was obtained after subtraction of the initial rate of NADPH consumption in absence of substrate. For the determination of coupling efficiency (CE), selectivity and total turnover number (TTN): In a cuvette, stirred at 900 rpm, 100 mM pH 8.0 potassium phosphate was supplemented with 0.2 mM testosterone and 0.24 mM NADPH. Reaction was started with addition of 100 nM P450_BM3_ enzyme with a final volume of 1 mL and it was monitored until NADPH depletion was constant (completion of the reaction). Afterwards, the reaction mixture was immediately transferred into 96 MTPs and frozen at -20°C. Reaction mixtures of 600 μL were taken and mixed with ethyl acetate (2 x 150 μL) with the liquid handling arm of the Tecan robot platform (dispensing speed, 600 μL/s) The organic phase was extracted using the Te-MO multi-pipette option, and transferred to 500 μL MTPs (Nunc, Roskilde, Denmark). The solvent was dried overnight, and the next day the steroid was resuspended in 150 μL acetonitrile and passed through a PTSF 96-well plate filter to remove particles (Pall, VWR, Germany) into a new 500 μL MTP (Nunc). The MTPs were stored at 4°C prior to screening. The kinetic parameters are shown in Supplementary Table 1.

