## Supplementary Information for "Pervasive cooperative mutational effects on multiple catalytic enzyme traits emerge via long-range conformational dynamics"

### Supplementary Notes

**Supplementary Note 1:** Kinetics of purified enzymes.

Due to the multicomponent reaction (NADPH, $O_2$, [S], [P]), the P450 non-Michaelis Menten kinetics[1] as well as the problem that kcat/Km does not consider the effects of changing substrate concentration and product inhibition during the biocatalytic reaction (which may lead to unreal expectations on enzyme performance and erroneous selection of the most adequate biocatalyst[2]), we only determined the following parameters to assess the catalytic efficiency of the $P450_{BM3}$ variants:

- Selectivity (2β hydroxytestosterone) based on %HPLC (values can be converted to KJ per mol[3])
- Substrate (testosterone) conversion based on %HPLC
- Product formation rate (PFR) in µmol products · µmol $enzyme^{-1}$ · $min^{-1}$
- NADPH consumption rate (NCR) in µmol NADPH · µmol $enzyme^{-1}$ · $min^{-1}$
- Coupling efficiency (CE) in percentage (%)
- Total turnover number (TTN) in µmol products · µmol $enzyme^{-1}$
- Total turnover frequency (TTF) in µmol products · µmol $enzyme^{-1}$ · $min^{-1}$

**Supplementary Note 2:** Quantum mechanic calculation.

The P450 protein-bound porphyrin complex coordinated with iron and containing an Fe=O moiety is known as compound I (Cpd I). In our computational model for Cpd I, substituents at the periphery of the porphyrin moiety were replaced with hydrogen atoms. Additionally, the axial Fe−SCys bond in Cpd I was modeled as a Fe−$SCH_3$ bond. Hydroxylation by P450 consists in the hydrogen atom abstraction from the C–H bond of the substrate by CpdI (which is the rate limiting step), followed by radical rebound mechanism. In agreement with previous work[4] the quartet C–H abstraction transition-states have been calculated to be consistently lower in energy than the doublet spin states. Hence, all the reported activation free-energies ($\Delta G^{\ddagger}$) correspond to C–H abstraction on the quartet potential energy surface.

A comprehensive computational DFT study was then performed with the Gaussian09(D.01)[5] program package using the B3LYP[6,7] hybrid functional under the unrestricted formalism (except for **1**, restricted singlet) and LANL2DZ pseudopotential with associated basis set[8] to describe the Fe atom and relativistic effects. All the other atoms were treated with a 6-31g(d) basis set during geometry optimization and subsequent frequency calculation, whereas 6-311(d,p) basis set was employed for final energies single point calculations. In all the calculations, solvent effects were taken into account using the Polarizable Continuum Model[9] (PCM) and dispersion energy correction was treated under the Grimme's $D_3$ formalism[10]. Supplementary Table 11 summarizes the energies, thermal corrections and free energies of the QM structures calculated for H-abstraction catalyzed by haem.

**Supplementary Note 3:** Molecular Dynamics simulations.

Long-timescale conventional Molecular Dynamics simulations (cMD) in explicit water were performed using AMBER 16 package[11] in our in-house GPU cluster Galatea. Substrate testosterone (**1**) parameters for the MD simulations were generated within the antechamber module of AMBER 16 using the general AMBER force field (GAFF),[12] with partial charges set to fit the electrostatic potential generated at the HF/6-31G(d) level by the restrained electrostatic potential (RESP) model.[13] The charges were calculated according to the Merz-Singh-Kollman scheme[14,15] using Gaussian 09(D.01).[5] Parameters for the haem resting state, Cpd I and axial Cys were taken from elsewere.[16] Amino acid protonation states were predicted using the H++ server (http://biophysics.cs.vt.edu/H++).[17] Then, the enzyme was solvated in a pre-equilibrated truncated hexagonal box with a 10-Å buffer of TIP3P[18] water molecules using the AMBER16 leap module, resulting in the addition of ~13,000 solvent molecules. The systems were neutralized by addition of explicit counterions ($Na^+$ and $Cl^-$). All subsequent calculations were done using the widely tested Lindorff-Larsen modification of the Amber 99 force field (ff99SBildn).[19] For all the mutants, the apo structures used were generated from the P450-BM3 original variant (PDB: 1FAG)[20], by removal of the palmitoleic acid bound to the protein and the introduction of the corresponding mutations using the RosettaBackrub web-server (https://kortemmeweb.ucsf.edu/backrub).[21–23] Substrate-bound structures were produced using the former mutants generated with RosettaBackrub as template, by placing the testosterone (**1**) molecule in the position originally occupied by the palmitoleic acid. Pose 2 and pose 15 were obtained by orienting **1** with its C2 or C15 atom towards the haem-moiety, respectively. The stability of all the aforementioned apo and substrate-bound structures was tested by MD simulations. A two-stage geometry optimization approach was performed. The first stage minimizes the positions of solvent molecules and ions imposing positional restraints on solute by a harmonic potential with a force constant of 500 kcal $mol^{-1}$ $Å^{-2}$, and the second stage is an unrestrained minimization of all the atoms in the simulation cell. The systems are gently heated using six 50-ps steps, incrementing the temperature 50 K each step (0–300 K) under constant volume and periodic boundary conditions. Water molecules were treated with the SHAKE algorithm such that the angle between the hydrogen atoms is kept fixed. Long-range electrostatic effects were modeled using the particle-mesh-Ewald method.[24] An 8-Å cutoff was applied to Lennard-Jones and electrostatic interactions. Harmonic restraints of 10 kcal/mol were applied to the solute, and the Langevin equilibration scheme was used to control and equalize the temperature. The time step was kept at 1 fs during the heating stages, allowing potential inhomogeneities to self-adjust. Each system was then equilibrated without restrains for 2 ns with a 2-fs timestep at a constant pressure of 1 atm and temperature of 300 K. After the systems were equilibrated in the NPT ensemble, for both apo and substrate-bound structures, three independent 600 ns MD simulations (i.e. 1.8 μs accumulated) were performed under the NVT ensemble and periodic-boundary conditions using our Galatea cluster (composed by 178 GTX1080 GPUs). With Galatea, simulations for these systems were performed at a speed of ca. 100 ns/day.

**Supplementary Note 4:** Accelerated Molecular Dynamic Simulations.

Long-timescale accelerated Molecular Dynamics simulations (aMD)[25–27] have been used to explore the binding trajectory of **1** in the different mutants. To allow the substrate to diffuse freely until being spontaneously recognized by the enzyme surface and finally getting in the access channel, we started with four different molecules of **1** randomly placed in the bulk solvent, performing three independent replicas, each one of 250 ns unconstrained cMD simulations (*vide supra*), from which the acceleration parameters were determined, followed by 750 ns of dual-boost aMD.

aMD enhances the conformational sampling of biomolecules, by adding a non-negative boost potential to the system when the system potential is lower than a reference energy:

$$V^*(r) = V(r), \qquad V(r) \geq E, \qquad (1)$$

$$V^*(r) = V(r) + \Delta V(r), \qquad V(r) < E \qquad (2)$$

where $V(r)$ is the original potential, $E$ is the reference energy, and $V^*(r)$ is the modified potential. In the simplest form, the boost potential, $\Delta V(r)$ is given by:

$$\Delta V(r) = \frac{[E-V(r)]^2}{\alpha+E-V(r)} \qquad (3)$$

where $\alpha$ is the acceleration factor. As the acceleration factor α decreases, the energy surface is flattened more and biomolecular transitions between the low-energy states are increased. Here a total boost potential is applied to all atoms in the system in addition to a more aggressive dihedral boost, i.e., ($E_{dihed}$, $\alpha_{dihed}$; $E_{total}$, $\alpha_{total}$), within the dual-boost aMD approach. The acceleration parameters used in this work, are the following:

$$E_{dihed} = V_{dihed\_avg} + 2.0 \times N_{res},\ \alpha_{dihed} = 2.0 \times \mathrm{N_{res}}/5, \qquad (4)$$

$$E_{total} = V_{total\_avg} + 0.16 \times N_{atoms},\ \alpha_{total} = 0.16 \times N_{atoms} \qquad (5)$$

where $N_{res}$ is the number of protein residues, $N_{atoms}$ is the total number of atoms, and $V_{dihed\_avg}$ and $V_{total\_avg}$ are the average dihedral and total potential energies calculated from 250 ns cMD simulations, respectively.

**Supplementary Note 5:** Additivity terms and equations.

Mathematically, the addition between A and B can be determined according to the trait in quest.

In the case of stereoselectivity or enantioselectivty[3], the free energy of interaction ($\Delta G^{\neq}_{AB}$) between two mutations (or sets of mutations) *A* and *B* can be calculated using equation (6):

$$\Delta \mathrm{G}^{\neq}_{AB} = \Delta\Delta \mathrm{G}^{\neq}_{exp} - (\Delta\Delta \mathrm{G}^{\neq}_{A} + \Delta\Delta \mathrm{G}^{\neq}_{B}) \qquad (6)$$

Where $\Delta\Delta \mathrm{G}^{\neq}_{exp}$ is the difference in activation energy with respect to regioselectivity (**2** versus **3** formation and all the rest of the stereo- and regioisomers) experimentally obtained for the binary combination, while $\Delta\Delta \mathrm{G}^{\neq}_{A}$ and $\Delta\Delta \mathrm{G}^{\neq}_{B}$ are the experimental energies obtained for each mutant (or sets of mutations) independently.

In the case of conversion (or any other trait), herein we introduce equation (7):

$$\Delta \mathrm{C}^{\%}_{AB} = \Delta \mathrm{C}^{\%}_{exp} - \left(\Delta \mathrm{C}^{\%}_{A} + \Delta \mathrm{C}^{\%}_{B}\right) \qquad (7)$$

Where $\Delta \mathrm{C}^{\%}_{exp}$ is the difference in conversion for **2** and the parent mutant F87A experimentally obtained for the binary combination, while $\Delta\%C_A$ and $\Delta\%C_B$ are the differences in the experimental conversion values between each mutant (or sets of mutations) separately and that of the parent mutant F87A (28 ± 1%): The conversion values are not logarithmic (in contrast to the selectivity ones) and thus the value of the parental mutant F87A has to be subtracted from the single-point mutants and the double mutant.

In any case, additivity for selectivity (conversion) occurs when $\Delta \mathrm{G}^{\neq}_{AB} = 0\ (\Delta \mathrm{C}^{\%}_{AB} = 0)$, while positive (+ME) and negative (-ME) magnitude epistasis correspondingly pertains when $\Delta \mathrm{G}^{\neq}_{AB} > 0\ (\Delta \mathrm{C}^{\%}_{AB} > 0)$ and $\Delta \mathrm{G}^{\neq}_{AB} < 0\ (\Delta \mathrm{C}^{\%}_{AB} < 0)$. Positive (+SE) and negative (-SE) sign epistasis occurs when $\Delta \mathrm{G}^{\neq}_{AB} > 0\ (\Delta \mathrm{C}^{\%}_{AB} > 0)$ if $A < 0\ or\ B < 0$ and $\Delta \mathrm{G}^{\neq}_{AB} < 0\ (\Delta \mathrm{C}^{\%}_{AB} < 0)$ if $A > 0\ or\ B > 0$, respectively. Finally, positive (+RSE) and negative (-RSE) reciprocal sign epistasis occurs when $\Delta \mathrm{G}^{\neq}_{AB} > 0\ (\Delta \mathrm{C}^{\%}_{AB} > 0)$ if $A < 0\ and\ B < 0$ or $\Delta \mathrm{G}^{\neq}_{AB} < 0\ (\Delta \mathrm{C}^{\%}_{AB} < 0)$ if $A > 0\ and\ B > 0$, respectively[28].

To exemplify the various cases of additivity/non-addivity, if the desired fitness trait is **2-beta hydroxytestosterone selectivity**, the following theoretical examples are given:

1. **Additivity:** Mutant A is 70% 2-beta sel. (2.1 KJ/mol). Mutant B is 79% 2-beta sel. (3.3 KJ/mol). Mutant AB is 90% 2-beta sel. (5.4 KJ/mol).

2. **Positive Magnitue Epistasis (+ME):** Mutant A is 70% 2-beta sel. (2.1 KJ/mol). Mutant B is 63% 2-beta sel. (1.3 KJ/mol). Mutant AB is 90% 2-beta sel. (5.4 KJ/mol).

3. **Positive Sign Epistasis (+SE):** Mutant A is 70% 2-beta sel. (2.1 KJ/mol). Mutant B is 63% 15-beta sel. (-1.3 KJ/mol). Mutant AB is 90% 2-beta sel. (5.4 KJ/mol).

4. **Positive Reciprocal Sign Epistasis (+RSE):** Mutant A is 70% 15-beta sel. (-2.1 KJ/mol). Mutant B is 63% 15-beta sel. (-1.3 KJ/mol). Mutant AB is 90% 2-beta sel. (5.4 KJ/mol).

5. **Negative Magnitue Epistasis (-ME):** Mutant A is 70% 2-beta sel. (2.1 KJ/mol). Mutant B is 63% 2-beta sel. (1.3 KJ/mol). Mutant AB is 60% 2-beta sel. (1.0 KJ/mol).

6. **Negative Sign Epistasis (-SE):** Mutant A is 70% 2-beta sel. (2.1 KJ/mol). Mutant B is 63% 15-beta sel. (-1.3 KJ/mol). Mutant AB is 90% 15-beta sel. (-5.4 KJ/mol).

7. **Negative Reciprocal Sign Epistasis (-RSE):** Mutant A is 70% 2-beta sel. (2.1 KJ/mol). Mutant B is 63% 2-beta sel. (1.3 KJ/mol). Mutant AB is 90% 15-beta sel. (-5.4 KJ/mol).

Using equations (6) and (7), the degree of interaction (if any) between two mutations (or sets of mutations) can be calculated.

All the calculations for the double mutants were performed as exemplified above for the binary combination of two single mutations (A+B) for **2** (Entry 1-6, Supplementary Tables 2-8). The additivity analyses were also performed for the ternary (i+j+k) combination of **2** (Entry 7, Supplementary Tables 2-8).

**Supplementary Note 6:** Automated calculation of additivity using Python 3.7.

The present study yielded a consequent amount of data with different equations (Supplementary Note 5). It was decided to automatize the data analysis process to gain time and reliability. To do so, a python 3.7 code was written with the Spyder interface. The code is deposited as https://github.com/matteoferla/Epistasis_Calculator and is divided in two parts:

The first part requires the user's input on different variables. The user indicates the number of mutations and the names. The user also needs to choose the right algorithm. There are two algorithms: Selectivity (S) or Conversion (C), which will have an impact for the final output of the program. The latter can be used for any parameter, except enantioselectivity. The output (excel tables) names also need to be given.

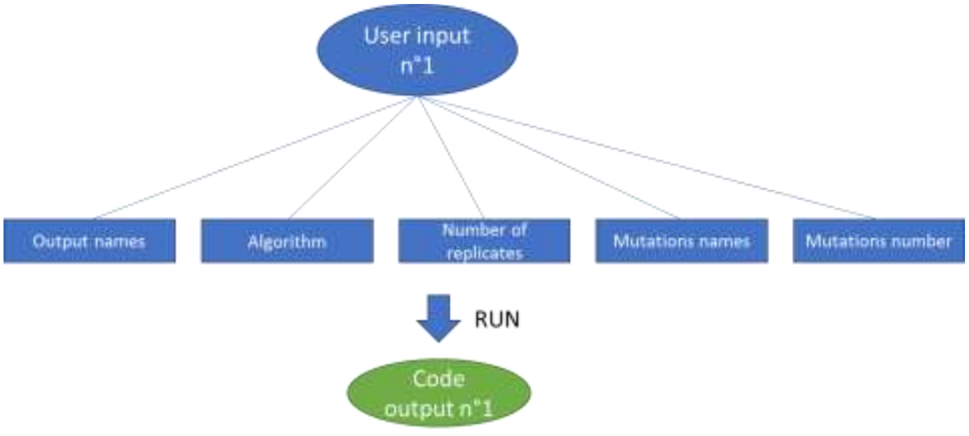

| | Mutation 1 | Mutation 2 | Mutation 3 | Replicate n°1 | Replicate n°2 | Replicate n°3 |
|---|---|---|---|---|---|---|
| Mutant 1 | - | - | - | X | X | X |
| Mutant 2 | - | - | + | X | X | X |
| Mutant 3 | - | + | - | X | X | X |
| Mutant 4 | + | - | - | X | X | X |
| Mutant 5 | + | + | - | X | X | X |
| Mutant 6 | + | - | + | X | X | X |
| Mutant 7 | - | + | + | X | X | X |
| Mutant 8 | + | + | + | X | X | X |

In the figure above, the user is asked to enter various variables. The code uses the variables to build its first output. In this table, the user will be able to enter data and proceed. In the example above, three mutations and three replicates were used. Their names were not specified for clarity. "X" filled cells are emplacement that the user needs to put its data in in order to proceed.

The program uses all these variables to build its first output which is a mutant table. The mutants are represented with signs "+" and "-" for each mutation. A "+" means that the mutant has the associated mutation. The program uses a randomized algorithm to generates all the possible mutants. Each time a new mutant is assembled, it is introduced in the table and if it is already present, the program discards it. The number of mutants is related to the number of mutations. For three mutations, eight mutants can be obtained, sixteen for four mutations. Because of the randomization, the first output varies for every use.

Once the user has entered the data in the first output, the program is used again on this table to yield the second final output. Using the different mutants, the program creates different combinations of mutants to obtain other mutants. For example, mutation 5 above is the result of the combination of mutations 3 and 4. Mutant 8 can be obtained by using either three different mutant pair combinations (6 and 3, 5 and 2, or 7 and 4) or a combination of three single mutations (2, 3 and 4). For a three mutations analysis, the number of combinations is seven, but it reaches thirty-six combinations when using four mutations.

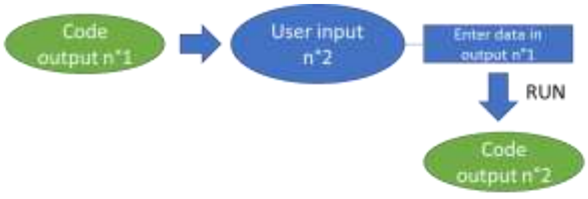

| | Mutation 1 | Mutation 2 | Mutation 3 | Combinations | Experimental average | Experimental standard deviation | Theoretical average | Theoretical standard deviation | Exp.avg - Theor.avg | Epistasis type |
|---|---|---|---|---|---|---|---|---|---|---|
| Combination n°1 | | - | + | {2, 3} | 79,444162 | 2,411977329 | 33,80111833 | 1,122567494 | 45,64308367 | "+ SE" |
| Combination n°2 | + | + | - | {3, 4} | 29,74640567 | 0,575299405 | 40,11042533 | 1,8107601 | -10,36401967 | "- SE" |
| Combination n°3 | + | - | + | {2, 4} | 26,51919833 | 1,337062731 | 45,412603 | 1,130403143 | -18,89340467 | "- ME" |
| Combination n°4 | + | + | + | {2, 5} | 68,412413 | 4,422365667 | 30,40450067 | 2,025752938 | 38,00791233 | "+ SE" |
| Combination n°5 | + | + | + | {4, 7} | 68,412413 | 4,422365667 | 86,411564 | 3,449793851 | -17,999151 | "- ME" |
| Combination n°6 | + | + | + | {3, 6} | 68,412413 | 4,422365667 | 21,87511567 | 2,009929371 | 46,53729733 | "+ RSE" |
| Combination n°7 | + | + | + | {2, 3, 4} | 68,412413 | 4,422365667 | 40,76852033 | 0,213766362 | 27,64389267 | "+ SE" |

In the second part of the program, the code assembles different combinations and attribute them different values based on the user's values. The program uses equations (6) or (7) given above (Supplementary Note 5) to give the results: Epistasis type. + = positive; - = negative; SE = "Sign Epistasis"; "ME" = Magnitude epistasis"; "RSE" = "Reciprocal Sign Epistasis".

After assembling the combinations, the program retrieves the values of their associated mutants and builds up "experimental" values. Each mutant of the first output is being given an average and standard deviation corresponding to the user's values. The program then assembles these values to yield "theoretical values". How the program uses these values is dictated by the "Selectivity" or "Conversion" algorithm. By comparing the theoretical values to the experimental ones, the program obtains a "score" (see Exp.avg – Theor.avg in previous figure). This score is what defines the nature of the epistatic interaction based on the equations (1) and (2) and the program gives its final result. The final output is a table containing theoretical and experimental addition as well as type and degree of epistatic interaction, if available. The server can be used to calculate any fitness value for any combination in protein evolution.

**Supplementary Note 7:** Pathway accessibility for multiple catalytic parameters.

The highest absolute value for each parameter in Fig. 2 of the main text was normalized to 100 % (mutant III shows the highest value for all parameters). Using the data obtained from the additivity equations (Supplementary Tables 2-8), we then calculated the theoretical values for each binary combination (indicated in empty bars with stripes) and compared it to the experimental data for each mutant (solid bars) in each pathway. See Supplementary Figure 11.

From the 6 possible pathways, two are non-accessible (33 %) because no improvements in fitness are observed. Fitness corresponds to the 7 different parameters of Supplementary Figure 11. Below, a and f exemplify an inaccessible and accessible pathway, respectively.

In the first step, the introduction of mutation R47I shows no epistatic effects for selectivity and coupling efficiency, and slightly synergistic effects for the remaining parameters; however, the pathway is not accessible because no catalytic improvement was obtained compared to the parent enzyme (a). Conversely, introduction of mutation Y51I results in synergistic effects on all parameters (particularly strong for selectivity and CE by >2-fold), but negative effects on conversion and no effects on NADPH consumption rate (f).

In the second step, combining mutation T49I with R47I (II-) provokes cooperative effects on selectivity and coupling efficiency without affecting conversion but antagonistic effects on TTN, TTF, PFR and NADPH consumption rate (a). In contrast, combining the same mutation T49I with Y51I (-II) results in synergistic effects on all parameters by 2-3-fold except for NADPH consumption rate, which remains almost unchanged (f).

Finally, introducing mutation Y51I into mutant II- results in pronounced cooperative effects by ~1.5- to 4-fold on all parameters excluding TTN (no epistatic effects) (a), whereas addition of mutation R47I to -II causes synergism in all parameters except TTN (i.e., additivity) (f).

This brief analysis shows that although all mutations are necessary for the emergence of positive epistatic effects, some pathways are more accessible than others depending on the fitness trait. For example, pathways starting with mutation Y51I show a high degree of selectivity (e, f), which is not observed in the other 4 pathways. Likewise, combining the previous mutation with T49I in two different pathways results in significant improvements in activity and in other parameters (d, f).

**Supplementary Note 8:** Conformational population analysis.

The conformational population analysis displayed in Figure 6c, 7a and Supplementary Fig 13 was obtained by applying the dimensionality reduction technique Principal Component Analysis (PCA) on the whole dataset of the apo trajectories (three replicas of 600 ns cMD for each mutant, i.e. an accumulated simulation time of 14.4 μs) considering the distances between all alpha carbons using the PyEMMA[29] software. The conformational population analysis obtained from PC1, PC2, and PC3 was able to successfully discriminate between 2- and 15-selective mutants. To further evaluate the physical meaning of the conformational changes involved in the first Principal Component (PC1) in Figure 6b,c, the PC1/PC3 space was used as it provided a better separation of 2- and 15-selective variants. The following procedure was followed: (i) the trajectory of the two mutant -I- (replica2) and III (replica3) were chosen as they exhibit the major variation in conformation along PC1, being located at its minimum and maximum values, respectively; (ii) the PCA dataset of each of the two mutant were clusterized into 200 clusters; (iii) the cluster located closer to the minimum of the conformational populational analysis was chosen as a representative structure for the overlay.

**Supplementary Note 9:** Shortest Path Map analysis.

The initial step of the Shortest Path Map (SPM) analysis generates a map based on the mean distances and correlation values computed along the MD simulation. For any residue of the protein a node is created and centered on its $C_\alpha$ if a mean distance shorter than 6 Å is explored with other residues along the simulation time. Subsequently, the length of the line connecting the nodes is drawn according to the correlation value of the corresponding residues ($d_{ij}$=-log $|C_{ij}|$). Larger correlation values (closer to |1|) will have shorter edge distances, while less correlated residue pairs (values closer to 0) will have longer edges distances. The Dijkstra algorithm is finally applied to identify the shortest path lengths. The algorithm goes through all the nodes of the graph and determines which is the shortest path to go from the first until the last protein residue. Hence, the method identifies the shorter edges of the graph (i.e. highly correlated), that are more frequently used for going through all residues of the protein (i.e. they are more central for the communication pathway). The SPM method is discussed in details in our recent work.[30]

**Supplementary Note 10:** Additional Packages.

The Kernel Density Estimation (KDE) plots in Fig. 3 and Supplementary Figures 6 and 7 were prepared monitoring the geometrical parameters (i.e. (haem)O···2/15C(TES) distance and (haem)O···H–2/15C(TES) angle) from cMD trajectories with **1** bound in pose 2 and pose 15 in the active site of the different mutants, using the Seaborn python library.[31] The principal component 2 (pc2) shown in Figures 4d, 6b and Supplementary Figure 12 was computed from the cumulated apo trajectory of the different mutants using the Principal Component Analysis (PCA) available in the Bio3D package.[32,33] The volumes of (i) the active site in mutants --- and III (Supplementary Figure 8) and (ii) the area of the access channel 2 in mutant -I-, III (Figure 7c) were calculated with the POVME2 software,[34] with a 1.0 Å grid spacing and 1.35 Å distance cutoff. In (i) the calculation was done on the most populated clusters of the joint apo trajectories of the two mutants, with a sphere of 6 Å as the including region, whereas in (ii) the structures used were those arising from the conformational population analysis (Supplementary Note 8, *vide supra*) with a sphere of 4 Å as the including region. All the images of the P450 mutants shown were done using the PyMOL software.[35]

#### Supplementary Figures

**Supplementary Figure 1:** Catalytic mechanism for aliphatic C-H hydroxylation catalyzed by P450s.

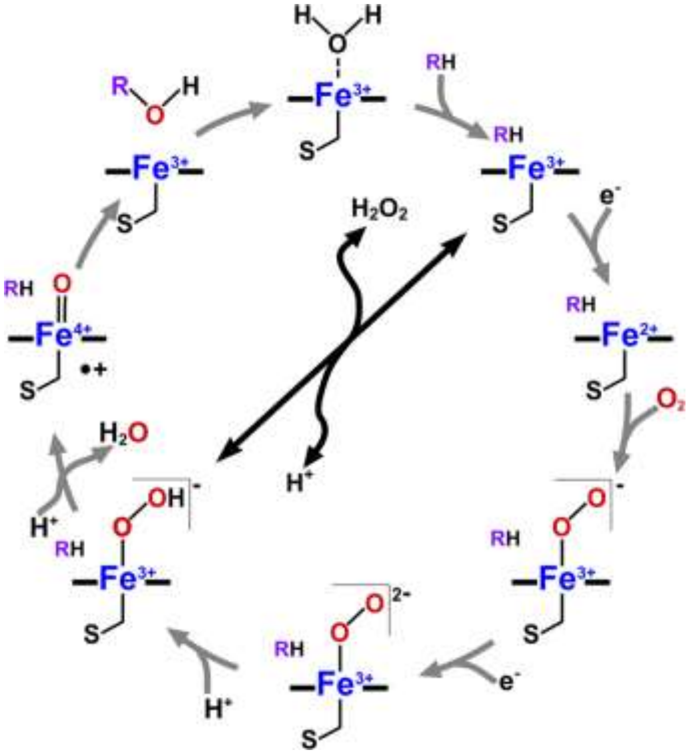

**Legend:** Figure taken from Ref. [36].

**Supplementary Figure 2:** SDS-PAGE analysis of purified mutants.

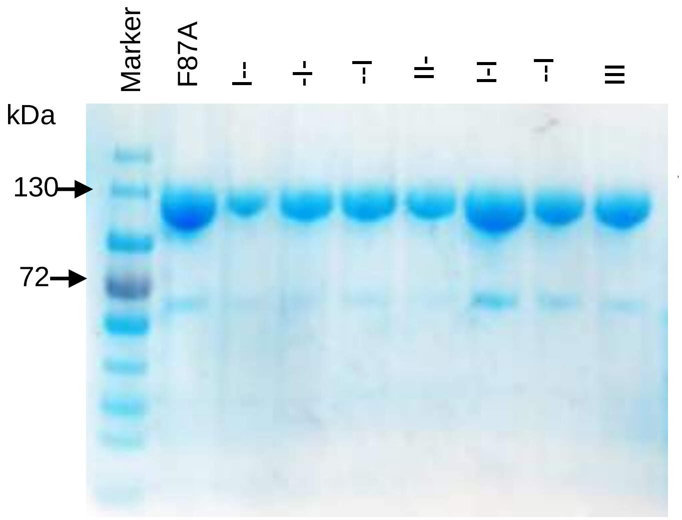

**Legend:** All N-terminal his-tagged proteins show purity above 80-90%.

**Supplementary Figure 3:** QM transition-states for the C–H abstraction in 1 from (a) C-2 (b) C-15.

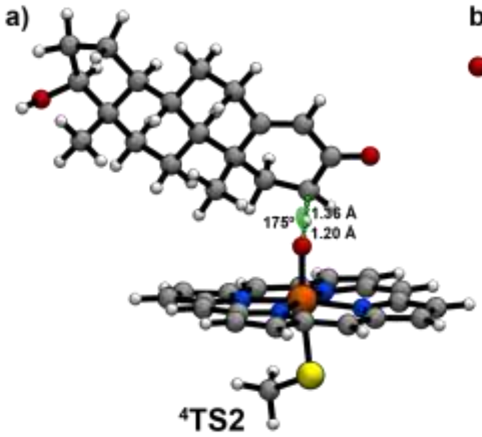

**Supplementary Figure 4:** Snapshot of **1** binding pose of during MD simulations in 15β-selective mutants.

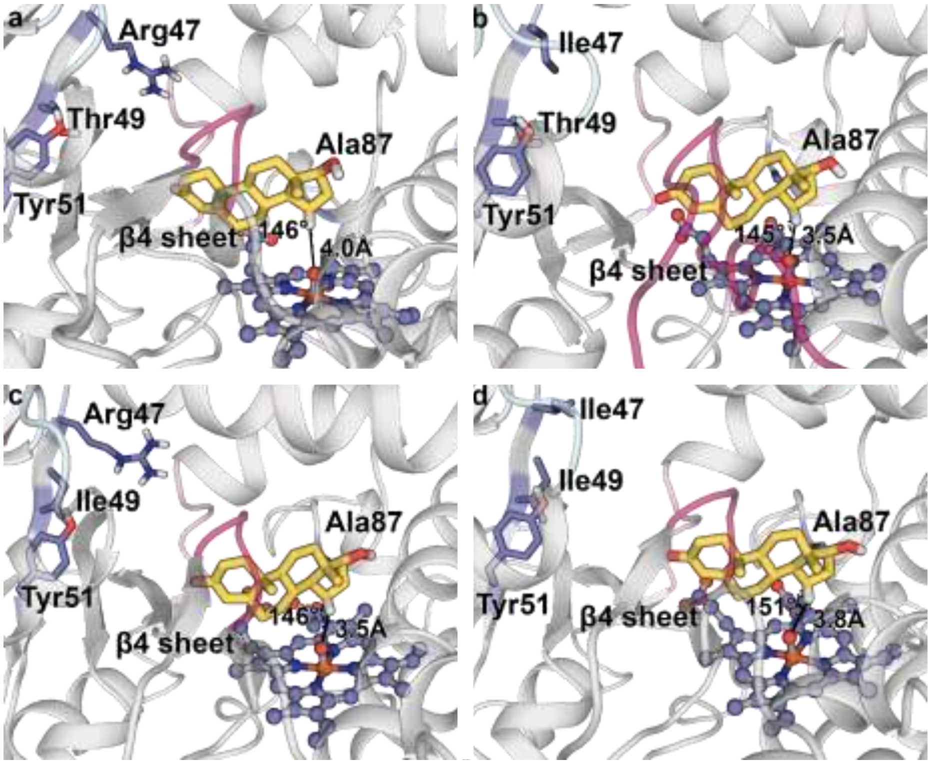

**Legend:** Mutants (a) ---; (b) I--; (c) -I- and (d) II- with distances measured between the oxygen atom of the Fe=O and the C15 atom of **1** and angles formed by O(Fe=O) – (**1**)-H(C15) – (**1**)-C(15).

**Supplementary Figure 5:** Snapshot of **1** binding pose during MD simulations in 2β-selective mutants.

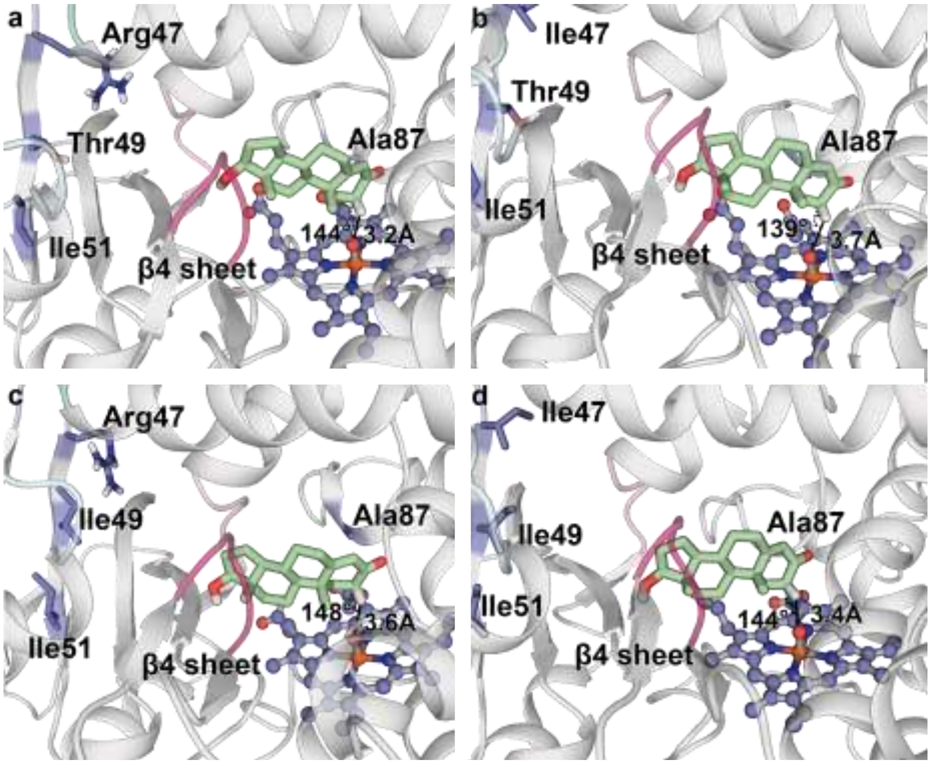

**Legend:** Mutants (a) I--; (b) I-I; (c) -II and (d) III with distances measured between the oxygen atom of the Fe=O and the C2 atom of **1** and angles formed by O(Fe=O) – (**1**)-H(C2) – (**1**)-C(2).

**Supplementary Figure 6.** KDE plot of the second replica dataset.

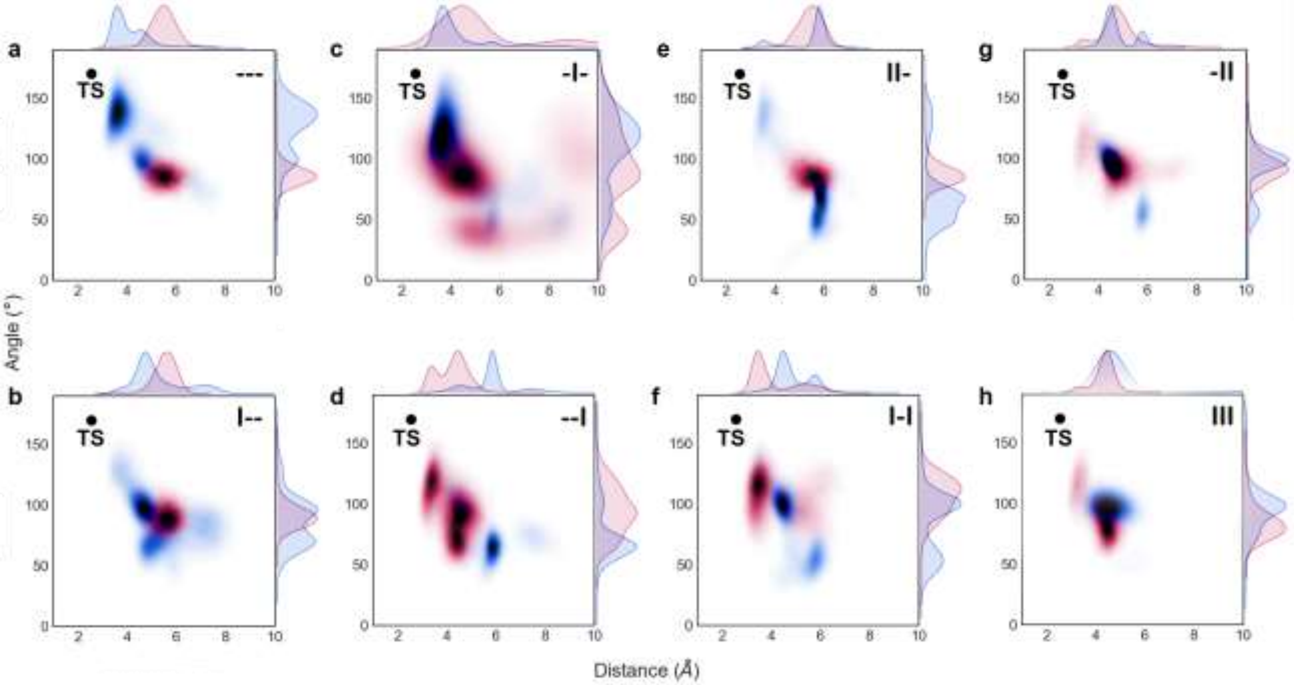

**Legend**: Distances determined between the oxygen atom of the Fe=O and the carbon atom (C2 or C15) of **1** (x-axis) and angles formed by O(Fe=O) – (**1**)-H(C2/15) – (**1**)-C(2/15) (y axis) from MD trajectories of mutants (a) ---, (b) I--, (c) -I-, (d) --I, (e) II-, (f) I-I, (g) -II and (h) III. Geometric parameters measured for carbon 2 (C2) and carbon 15 (C15) are shown in red and red blue, respectively. The ideal distance and angle for the TS (black dot) corresponds to the Density Functional Theory (DFT) optimized geometry for the C–H abstraction by Fe=O using a truncated computational model.

**Supplementary Figure 7.** KDE plot of the third replica dataset.

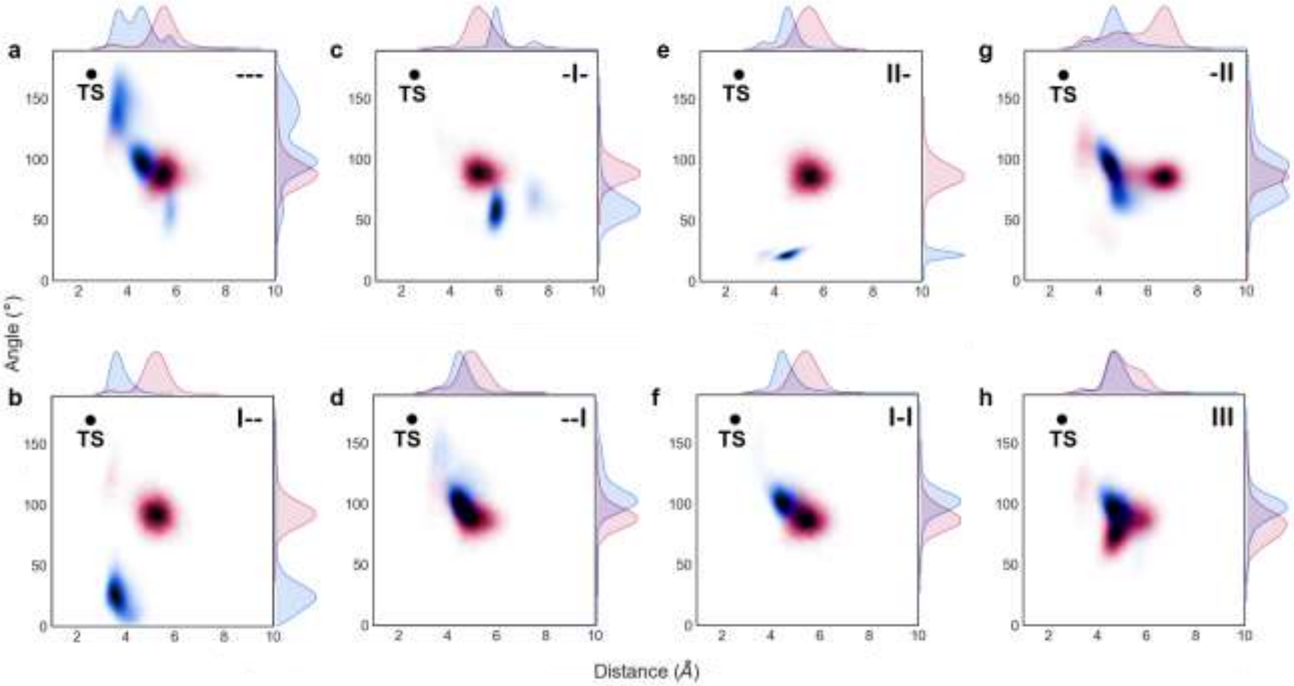

**Legend**: Distances determined between the oxygen atom of the Fe=O and the carbon atom (C2 or C15) of **1** (x-axis) and angles formed by O(Fe=O) – (**1**)-H(C2/15) – (**1**)-C(2/15) (y axis) from MD trajectories of mutants (a) ---, (b) I--, (c) -I-, (d) --I, (e) II-, (f) I-I, (g) -II and (h) III. Geometric parameters measured for carbon 2 (C2) and carbon 15 (C15) are shown in red and red blue, respectively. The ideal distance and angle for the TS (black dot) corresponds to the Density Functional Theory (DFT) optimized geometry for the C–H abstraction by Fe=O using a truncated computational model.

**Supplementary Figure 8:** Computed active site volume for mutants (a) (III) and (b) ---.

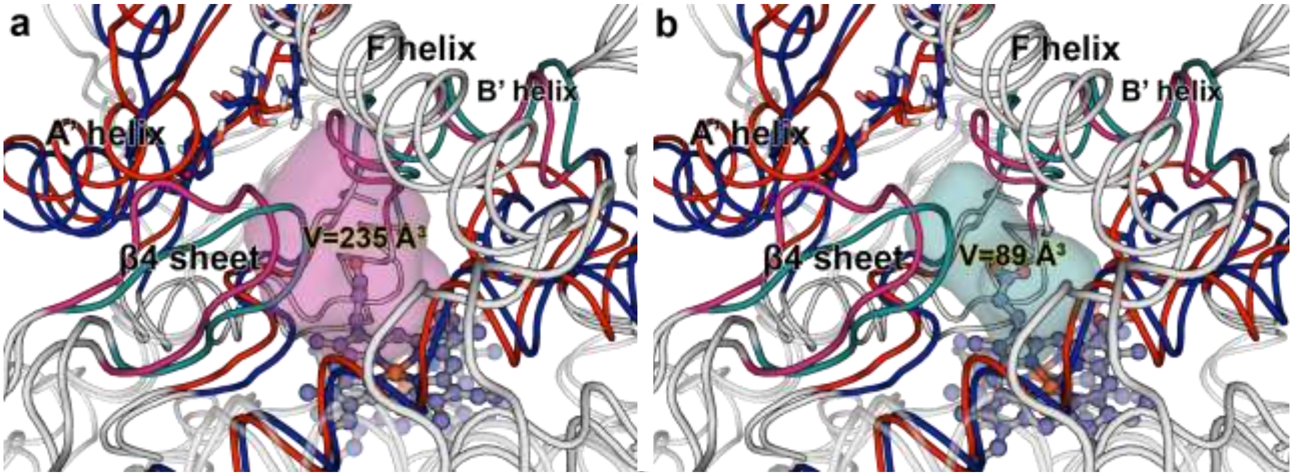

**Legend:** Mutant I-- is shown in blue, whereas the evolved mutant III is coloured in red. The F-G loop and the β1 sheet / active site volume are highlighted in teal and magenta colour for -I- and III, respectively.

**Supplementary Figure 9:** Final pose of **1** in the aMD binding trajectory of I--.

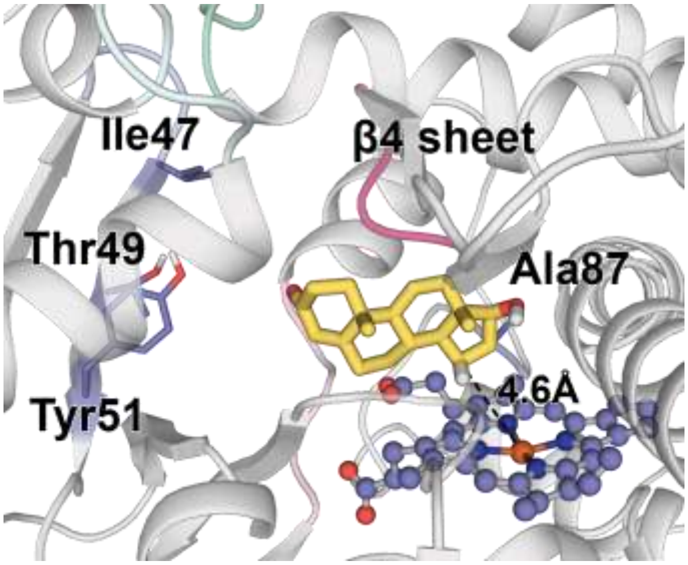

**Supplementary Figure 10:** (**1**)CO-Y51 polar interaction interaction.

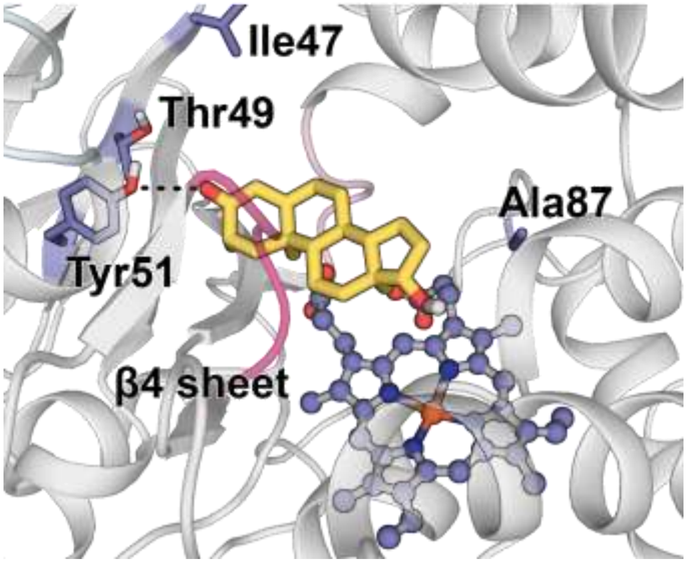

**Supplementary Figure 11:** Pathway accessibility for multiple enzymatic parameters.

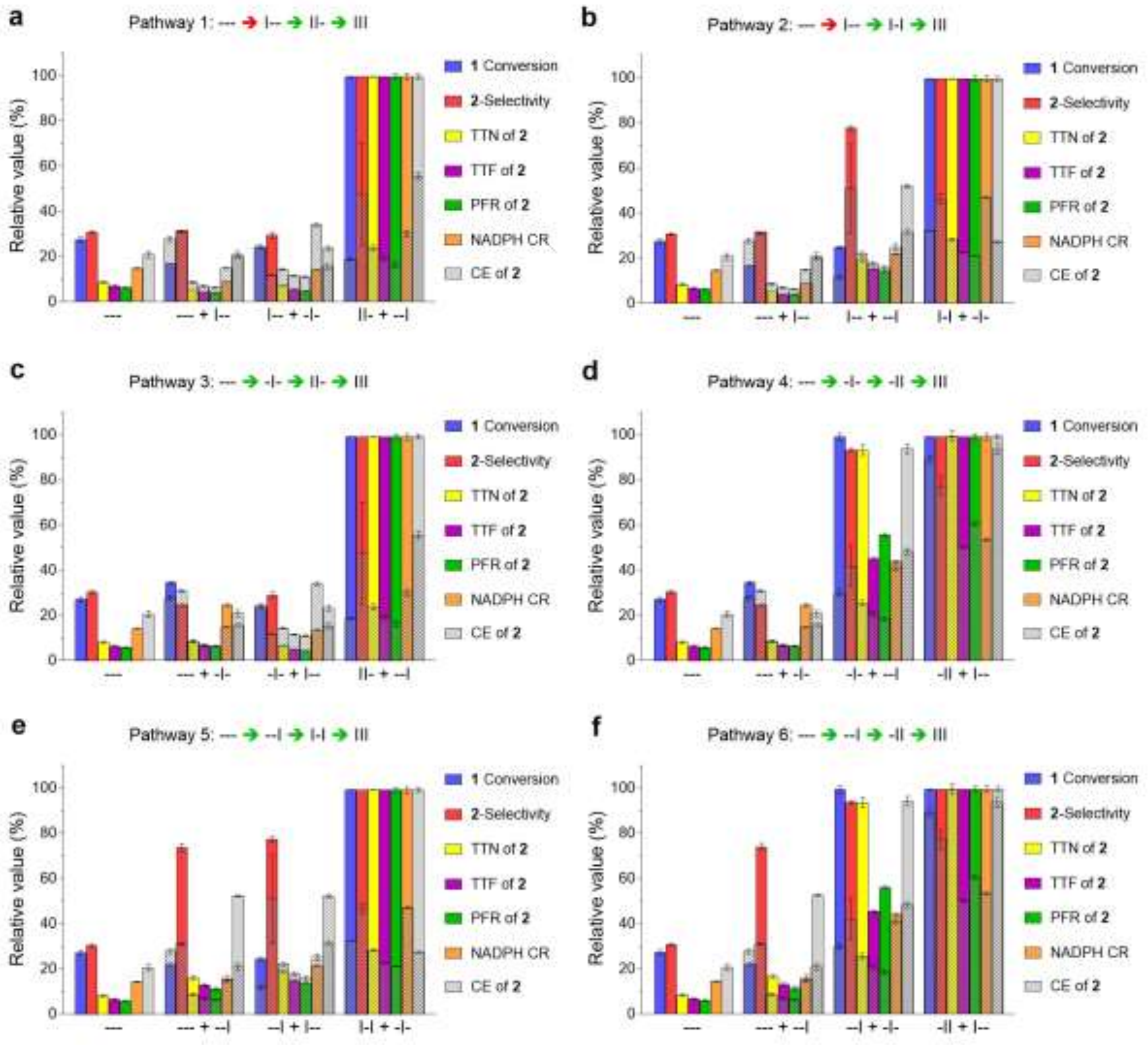

**Legend:** Stepwise multiparametric pathway analysis. Pathway 1 (a) or 2 (b) start with mutant F87A towards mutation R47I generating double mutant R47I/F87A. This step decreases all the parameters compared to the parent enzyme F87A (red arrow) but in the second and third step, the individual introduction of mutation T49I or Y51I and Y51I or T49I, respectively, improves all parameters. Pathway 3 (c) or 4 (d) begin with mutation T49I, while pathway 5 (a) or 6 (b) start with mutation Y51I. All these pathways show green arrows, indicating the absence of local minima. The experimental data for each mutant is shown in solid bars and normalized to 100% regarding the highest value. The expected theoretical values (using additivity equations) are indicated in empty bars with stripes. The data represent the average ± s.e.m. of two independent experiments (*n* = 2).

**Supplementary Figure 12:** PC Analysis (pc2) of apo-state mutated enzymes along the fitness landscape pathways.

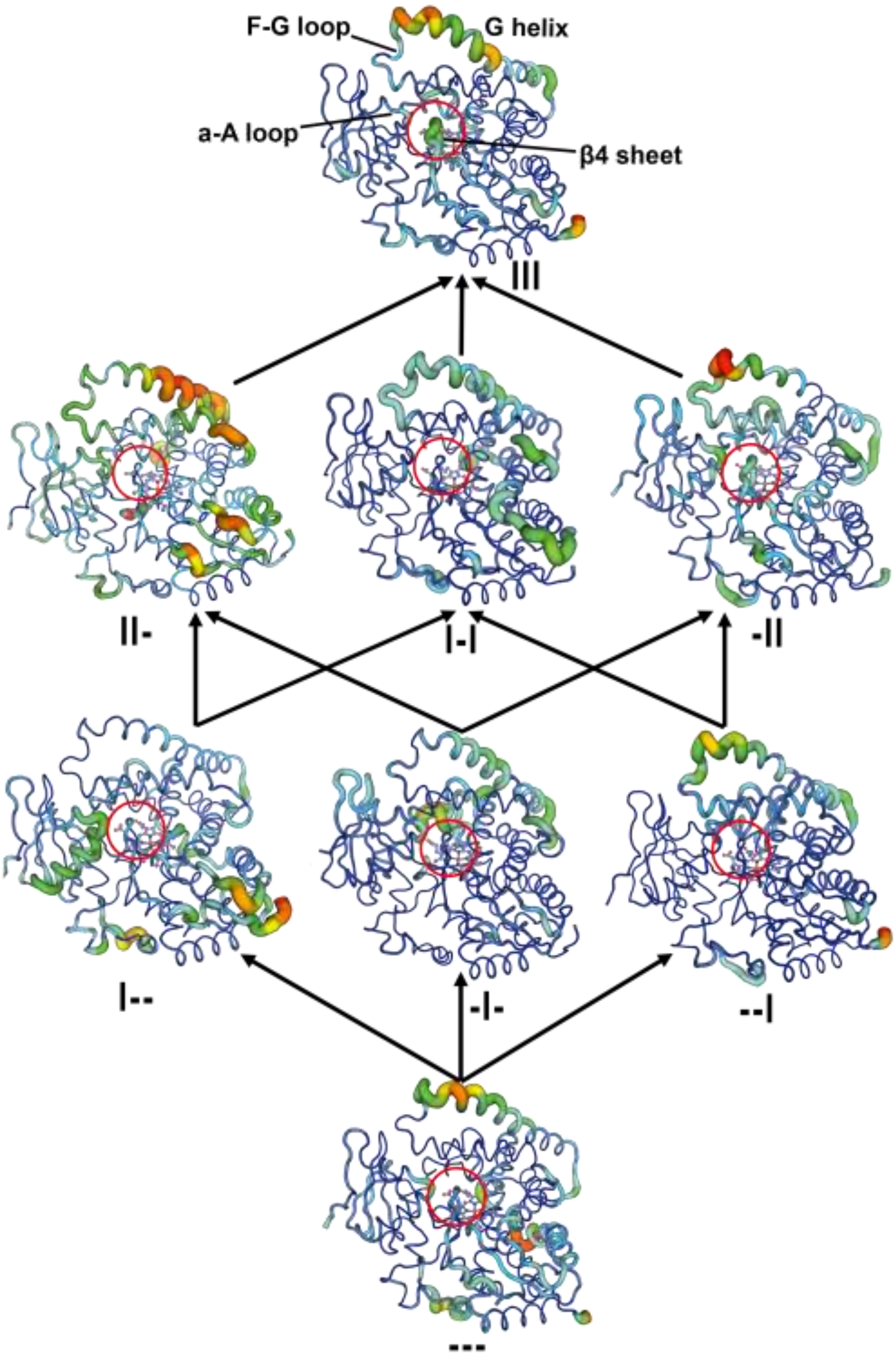

**Legend:** The thickness of the line is proportional to the motion and the colour scale varies from blue (minimum motion) to red (maximum motion).

**Supplementary Figure 13:** Conformational population analysis (pc1, pc2) of deconvoluted mutants.

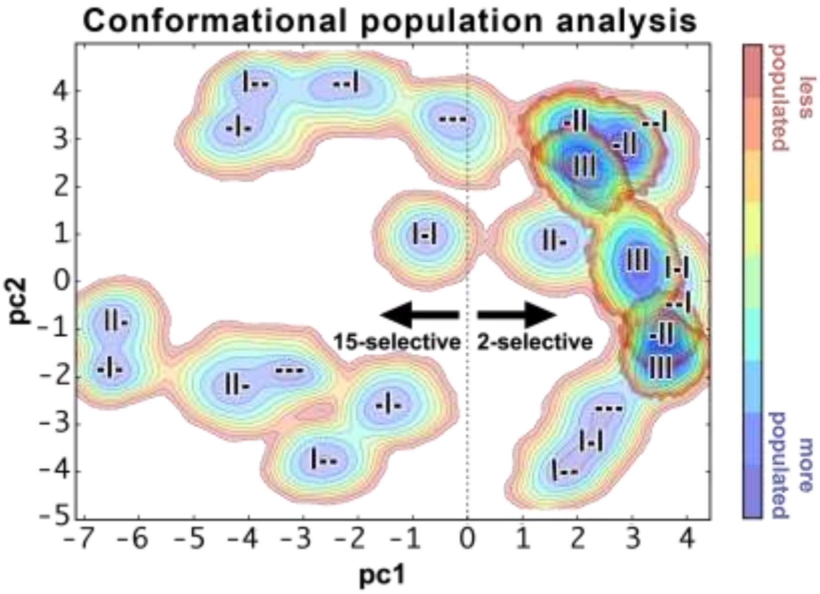

**Legend:** The free energies of 2β-selective mutants -II and III are highlighted.

### Supplementary Tables

#### **Supplementary Table 1:** Kinetic profiles of deconvoluted mutants.

| Entry | Mutant | Conc. (µM) | Initial rate (µmol / µmol x s) | 1 Conv. HPLC (%) | CE (%) | 2-Selectivity | | TTN (µmol/µmol) | TTF (µmol / µmol x s) |
|---|---|---|---|---|---|---|---|---|---|
| | | | | | | HPLC (%) | ΔΔG‡ (kJ per mol) | | |
| 1 | --- | 1.5 | 50 ± 4 | 10 ±1 | 25 ± 2 | 28 ± 1 | -2.323 ± 0.124 | 194 ± 11 | 11 ± 0.4 |
| 2 | I-- | 1.3 | 25 ± 6 | 6 ± 0 | 23 ± 0 | 29 ± 0 | -2.229 ± 0.051 | 120 ± 1 | 7 ± 0.1 |
| 3 | -I- | 0.9 | 115 ± 11 | 12 ± 0 | 24 ± 1 | 23 ± 1 | -3.015 ± 0.136 | 246 ± 6 | 11 ± 0.2 |
| 4 | --I | 1.9 | 90 ± 16 | 8 ± 0 | 26 ± 1 | 67 ± 2 | 1.802 ± 0.222 | 157 ± 8 | 21 ± 0.7 |
| 5 | II- | 1.5 | 54 ± 4 | 8 ± 0 | 30 ± 0 | 27 ± 2 | -2.494 ± 0.259 | 168 ± 3 | 9 ± 0.1 |
| 6 | I-I | 1.3 | 96 ± 6 | 9 ± 0 | 15 ± 0 | 71 ± 1 | 2.189 ± 0.119 | 175 ± 6 | 24 ± 0.6 |
| 7 | -II | 3.2 | 337 ± 6 | 35 ± 1 | 37 ± 1 | 85 ± 1 | 4.358 ± 0.192 | 700 ± 16 | 72 ± 1.2 |
| 8 | III | 2.0 | 588 ± 1 | 35 ± 0 | 37 ± 1 | 91 ± 0 | 5.594 ± 0.052 | 701 ± 1 | 158 ± 0.2 |

Mutant: (---) = F87A, (-) = absence of mutation(s) at position(s) R47, T49 and/or Y51, III = R47I/T49I/Y51I/F87A. Concentration of purified enzymes was determined by CO difference spectrum analysis. The reactions were performed using 100 nM enzyme, 0.2 mM testosterone (**1**) as substrate and 1% DMF as cosolvent and 0.24 mM NADPH in potassium phosphate (KPi) buffer (pH 8.0) at 25°C until NADPH depletion, which was measured at 340 nm. The initial rate of product formation vs. NADPH consumption in the presence of substrate is based on the first 20 s. The relationship between total product formed and NADPH consumption is given by the coupling efficiency (CE). The total turnover number (TTN) is calculated from the amount of product formed and enzyme concentration used. ΔΔG‡ (KJ/mol) = Ln (2β-hydroxytestosterone / all other regio- and -stereoisomers) x (298 K x 0.001 x 8.31441 J $mol^{-1}$ $K^{-1}$). Data represent the average ± s.e.m. of two independent experiments (*n* = 2).

#### **Supplementary Table 2:** Additivity calculations for substrate conversion.

| Combination # | R47I | T49I | Y51I | Mutations | Exp. Mean | Exp. SEM | Theor. Mean | Theor. SEM | Exp. - Theor. (Mean) | Epistasis type |
|---|---|---|---|---|---|---|---|---|---|---|
| 1 | + | + | - | (2, 3) | 8.4 | 0.1 | 8.6 | 0.2 | -0.2 | Additive |
| 2 | - | + | + | (3, 4) | 35.0 | 0.6 | 10.5 | 0.5 | 24.6 | + Sign epistasis |
| 3 | + | - | + | (2, 4) | 8.8 | 0.2 | 4.1 | 0.3 | 4.6 | + Reciprocal sign epistasis |
| 4 | + | + | + | (2, 5) | 35.1 | 0.0 | 31.3 | 0.6 | 3.8 | + Sign epistasis |
| 5 | + | + | + | (4, 7) | 35.1 | 0.0 | 6.5 | 0.2 | 28.5 | + Reciprocal sign epistasis |
| 6 | + | + | + | (3, 6) | 35.1 | 0.0 | 11.4 | 0.0 | 23.7 | + Sign epistasis |
| 7 | + | + | + | (2, 3, 4) | 35.1 | 0.0 | 6.7 | 0.5 | 28.3 | + Sign epistasis |

The signs +/- mean that the respective mutation is present/absent. Mutation 2: R47I; Mutation 3: T49I; Mutation 4: Y51I. Mutation 5: T49I/Y51I; Mutation 6: R47I/Y51I; Mutation 7: R47I/T49I. The values are calculated with original data from **Supplementary Table 1**.

#### **Supplementary Table 3:** Additivity calculations for selectivity.

| Combination # | R47I | T49I | Y51I | Mutations | Exp. Mean | Exp. SEM | Theor. Mean | Theor. SEM | Exp. - Theor. (Mean) | Epistasis type |
|---|---|---|---|---|---|---|---|---|---|---|
| 1 | + | + | - | (2, 3) | -2.5 | 0.2 | -5.2 | 0.08 | 2.8 | + Magnitude epistasis |
| 2 | - | + | + | (3, 4) | 4.4 | 0.1 | -1.2 | 0.26 | 5.6 | + Sign epistasis |
| 3 | + | - | + | (2, 4) | 2.2 | 0.1 | -0.4 | 0.17 | 2.6 | + Sign epistasis |
| 4 | + | + | + | (2, 5) | 5.6 | 0.0 | 2.1 | 0.12 | 3.5 | + Sign epistasis |
| 5 | + | + | + | (4, 7) | 5.6 | 0.0 | -0.7 | 0.33 | 6.3 | + Sign epistasis |
| 6 | + | + | + | (3, 6) | 5.6 | 0.0 | -0.8 | 0.04 | 6.4 | + Sign epistasis |
| 7 | + | + | + | (2, 3, 4) | 5.6 | 0.0 | -3.5 | 0.26 | 9.0 | + Sign epistasis |

The signs +/- mean that the respective mutation is present/absent. Mutation 2: R47I; Mutation 3: T49I; Mutation 4: Y51I. Mutation 5: T49I/Y51I; Mutation 6: R47I/Y51I; Mutation 7: R47I/T49I. The values are calculated with original data from **Supplementary Table 1**.

#### **Supplementary Table 4:** Additivity calculations for total turnover number (TTN).

| Combination # | R47I | T49I | Y51I | Mutations | Exp. Mean | Exp. SEM | Theor. Mean | Theor. SEM | Exp. - Theor. (Mean) | Epistasis type |
|---|---|---|---|---|---|---|---|---|---|---|
| 1 | + | + | - | (2, 3) | 45 | 2 | 91 | 3 | -46 | - Magnitude epistasis |
| 2 | - | + | + | (3, 4) | 598 | 14 | 162 | 8 | 435 | + Magnitude epistasis |
| 3 | + | - | + | (2, 4) | 124 | 5 | 140 | 6 | -16 | - Magnitude epistasis |
| 4 | + | + | + | (2, 5) | 635 | 1 | 632 | 14 | 3 | Additive |
| 5 | + | + | + | (4, 7) | 635 | 1 | 151 | 8 | 484 | + Magnitude epistasis |
| 6 | + | + | + | (3, 6) | 635 | 1 | 180 | 2 | 455 | + Magnitude epistasis |
| 7 | + | + | + | (2, 3, 4) | 635 | 1 | 197 | 9 | 438 | + Magnitude epistasis |

The signs +/- mean that the respective mutation is present/absent. Mutation 2: R47I; Mutation 3: T49I; Mutation 4: Y51I. Mutation 5: T49I/Y51I; Mutation 6: R47I/Y51I; Mutation 7: R47I/T49I. The values are calculated with original data from **Supplementary Table 1**.

#### **Supplementary Table 5:** Additivity calculations for total turnover frequency (TTF).

| Combination # | R47I | T49I | Y51I | Mutations | Exp. Mean | Exp. SEM | Theor. Mean | Theor. SEM | Exp. - Theor. (Mean) | Epistasis type |
|---|---|---|---|---|---|---|---|---|---|---|
| 1 | + | + | - | (2, 3) | 8.7 | 0.4 | 18.2 | 0.5 | -9.4 | - Magnitude epistasis |
| 2 | - | + | + | (3, 4) | 72.2 | 0.9 | 32.4 | 1.7 | 39.8 | + Magnitude epistasis |
| 3 | + | - | + | (2, 4) | 24.3 | 0.8 | 28.1 | 1.2 | -3.8 | - Magnitude epistasis |
| 4 | + | + | + | (2, 5) | 157.8 | 0.0 | 79.1 | 1.0 | 78.7 | + Magnitude epistasis |
| 5 | + | + | + | (4, 7) | 157.8 | 0.0 | 29.9 | 1.6 | 127.8 | + Magnitude epistasis |
| 6 | + | + | + | (3, 6) | 157.8 | 0.0 | 35.6 | 0.3 | 122.2 | + Magnitude epistasis |
| 7 | + | + | + | (2, 3, 4) | 157.8 | 0.0 | 39.3 | 1.7 | 118.4 | + Magnitude epistasis |

The signs +/- mean that the respective mutation is present/absent. Mutation 2: R47I; Mutation 3: T49I; Mutation 4: Y51I. Mutation 5: T49I/Y51I; Mutation 6: R47I/Y51I; Mutation 7: R47I/T49I. The values are calculated with original data from **Supplementary Table 1**.

#### **Supplementary Table 6:** Additivity calculations for product formation rate (PFR).

| Combination # | R47I | T49I | Y51I | Mutations | Exp. Mean | Exp. SEM | Theor. Mean | Theor. SEM | Exp. - Theor. (Mean) | Epistasis type |
|---|---|---|---|---|---|---|---|---|---|---|
| 1 | + | + | - | (2, 3) | 12.4 | 0.8 | 28.6 | 1.0 | -16.2 | - Magnitude epistasis |
| 2 | - | + | + | (3, 4) | 147.8 | 2.0 | 48.5 | 2.8 | 99.3 | + Magnitude epistasis |
| 3 | + | - | + | (2, 4) | 37.4 | 1.0 | 41.1 | 2.6 | -3.7 | - Magnitude epistasis |
| 4 | + | + | + | (2, 5) | 262.3 | 3.0 | 158.4 | 2.4 | 103.8 | + Magnitude epistasis |
| 5 | + | + | + | (4, 7) | 262.3 | 3.0 | 42.9 | 3.1 | 219.4 | + Magnitude epistasis |
| 6 | + | + | + | (3, 6) | 262.3 | 3.0 | 55.4 | 0.4 | 206.9 | + Magnitude epistasis |
| 7 | + | + | + | (2, 3, 4) | 262.3 | 3.0 | 59.1 | 3.2 | 203.2 | + Magnitude epistasis |

The signs +/- mean that the respective mutation is present/absent. Mutation 2: R47I; Mutation 3: T49I; Mutation 4: Y51I. Mutation 5: T49I/Y51I; Mutation 6: R47I/Y51I; Mutation 7: R47I/T49I. The values are calculated with original data from **Supplementary Table 1**.

#### **Supplementary Table 7:** Additivity calculations for NADPH consumption rate (NCR).

| Combination # | R47I | T49I | Y51I | Mutations | Exp. Mean | Exp. SEM | Theor. Mean | Theor. SEM | Exp. - Theor. (Mean) | Epistasis type |
|---|---|---|---|---|---|---|---|---|---|---|
| 1 | + | + | - | (2, 3) | 50.3 | 0.7 | 120.9 | 2.7 | -70.6 | - Magnitude epistasis |
| 2 | - | + | + | (3, 4) | 157.4 | 2.2 | 145.5 | 3.6 | 11.9 | + Magnitude epistasis |
| 3 | + | - | + | (2, 4) | 78.1 | 1.2 | 88.6 | 5.3 | -10.5 | - Magnitude epistasis |
| 4 | + | + | + | (2, 5) | 354.3 | 5.1 | 189.4 | 2.7 | 164.9 | + Magnitude epistasis |
| 5 | + | + | + | (4, 7) | 354.3 | 5.1 | 106.9 | 5.0 | 247.4 | + Magnitude epistasis |
| 6 | + | + | + | (3, 6) | 354.3 | 5.1 | 167.0 | 1.0 | 187.3 | + Magnitude epistasis |
| 7 | + | + | + | (2, 3, 4) | 354.3 | 5.1 | 177.5 | 3.1 | 176.8 | + Magnitude epistasis |

The signs +/- mean that the respective mutation is present/absent. Mutation 2: R47I; Mutation 3: T49I; Mutation 4: Y51I. Mutation 5: T49I/Y51I; Mutation 6: R47I/Y51I; Mutation 7: R47I/T49I. The values are calculated with original data from **Supplementary Table 1**.

**Supplementary Table 8:** Additivity calculations for coupling efficiency (CE).

| Combination # | R47I | T49I | Y51I | Mutations | Exp. Mean | Exp. SEM | Theor. Mean | Theor. SEM | Exp. - Theor. (Mean) | Epistasis type |
|---|---|---|---|---|---|---|---|---|---|---|
| 1 | + | + | - | (2, 3) | 7.9 | 0.4 | 5.2 | 0.4 | 2.8 | + Reciprocal sign epistasis |
| 2 | - | + | + | (3, 4) | 31.7 | 0.7 | 16.2 | 0.4 | 15.5 | + Sign epistasis |
| 3 | + | - | + | (2, 4) | 10.7 | 0.4 | 17.5 | 0.2 | -6.7 | - Sign epistasis |
| 4 | + | + | + | (2, 5) | 33.5 | 0.4 | 31.4 | 0.8 | 2.1 | + Sign epistasis |
| 5 | + | + | + | (4, 7) | 33.5 | 0.4 | 18.7 | 0.5 | 14.8 | + Magnitude epistasis |
| 6 | + | + | + | (3, 6) | 33.5 | 0.4 | 9.2 | 0.1 | 24.3 | + Sign epistasis |
| 7 | + | + | + | (2, 3, 4) | 33.5 | 0.4 | 15.9 | 0.5 | 17.6 | + Sign epistasis |

The signs +/- mean that the respective mutation is present/absent. Mutation 2: R47I; Mutation 3: T49I; Mutation 4: Y51I. Mutation 5: T49I/Y51I; Mutation 6: R47I/Y51I; Mutation 7: R47I/T49I. The values are calculated with original data from **Supplementary Table 1**.

**Supplementary Table 9:** Pathway accessibility analysis based on selectivity.

| Pathway | Mutation | | | | ΔΔG‡ (kJ per mol) | | | |
|---|---|---|---|---|---|---|---|---|
| | 0 | 1st | 2nd | 3rd | 0 | 1st | 2nd | 3rd |
| 1 | --- | I-- | II- | III | -2.323 | -2.229 | -2.494 | 5.594 |
| 2 | --- | I-- | I-I | III | -2.323 | -2.229 | 2.189 | 5.594 |
| 3 | --- | -I- | II- | III | -2.323 | -3.015 | -2.494 | 5.594 |
| 4 | --- | -I- | -II | III | -2.323 | -3.015 | 4.358 | 5.594 |
| 5 | --- | --I | I-I | III | -2.323 | 1.802 | 2.189 | 5.594 |
| 6 | --- | --I | -II | III | -2.323 | 1.802 | 4.358 | 5.594 |

A green or red pathway/value means that there is a respective increase or decrease of fitness in each step of the evolutionary process from parental mutant F87A (---) toward triple mutant III (R47I/T49I/Y51I/F87A). Values in orange indicate that there is neither increase nor decrease in fitness between the precedent and subsequent mutation (considering SEM), thus its pathway turns red. Conversely to red, green pathways are accessible to stepwise evolution. For clarity, SEM values are not shown but can be found in **Supplementary Table 1.**

**Supplementary Table 10:** Pathway accessibility analysis based on TTF.

| Pathway | Mutation | | | | Total Turnover Number (TTN) | | | |
|---|---|---|---|---|---|---|---|---|
| | 0 | 1st | 2nd | 3rd | 0 | 1st | 2nd | 3rd |
| 1 | --- | I-- | II- | III | 11 | 7 | 9 | 158 |
| 2 | --- | I-- | I-I | III | 11 | 7 | 24 | 158 |
| 3 | --- | -I- | II- | III | 11 | 11 | 9 | 158 |
| 4 | --- | -I- | -II | III | 11 | 11 | 72 | 158 |
| 5 | --- | --I | I-I | III | 11 | 21 | 24 | 158 |
| 6 | --- | --I | -II | III | 11 | 21 | 72 | 158 |

A green or red pathway/value means that there is a respective increase or decrease of fitness in each step of the evolutionary process from parental mutant F87A (---) toward triple mutant III (R47I/T49I/Y51I/F87A). Values in orange indicate that there is neither increase nor decrease in fitness between the precedent and subsequent mutation (considering SEM), thus its pathway turns red. Conversely to red, green pathways are accessible to stepwise evolution. For clarity, SEM values are not shown but can be found in **Supplementary Table 1.**

**Supplementary Table 11:** Energies, thermal corrections and free energies of the QM structures calculated for H-abstraction catalyzed by haem.

| Structure[a] | $E_{el}$[b] | $E_{el}$[c] | G correction[d] | G[e] | ΔG |
|---|---|---|---|---|---|
| | (Hartree)[f] | (Hartree)[f] | (Hartree/particle)[f] | (Hartree)[f] | (kcal/mol) |
| **$^{4}$cpdI** | -1625.181930 | -1625.492613 | 0.267701 | -1625.224912 | |
| **1** | -891.402149 | -891.636974 | 0.397895 | -891.239079 | |
| **$^{4}$RC[g]** | - 2516.584079 | -2517.129587 | 0.665596 | -2516.463991 | |
| **$^{4}$TS2** | -2516.560005 | -2517.123777 | 0.681106 | -2516.442671 | 13.4 |
| **$^{4}$TS15** | -2516.559394 | -2517.123139 | 0.680743 | -2516.442396 | 13.6 |

[a] All the calculation were performed in the quartet state under the unrestricted formalism exept for **1** (singlet, restricted).
[b] Enegy obtained from geometries optimized at the B3LYP-$D_3$ / 6-31G* + LANL2DZ(Fe) level of theory.
[c] Energy obtained from single point calculation at the B3LYP-$D_3$ / 6-311G** + LANL2DZ(Fe) level of theory on top of the optimized geometry.
[d] Thermal correction at 298.15 K obtained from frequency calculation on top the optimized geometry at the same level of theory.
[e] Energy calculated as $E_{el}$[c] + G correction.
[f] 1 Hartree = 627.5 kcal/mol.
[g] Energy and thermal correction calculated as $^{4}$cpdI + **1**.

#### Supplementary Videos

**Supplementary Video 1:** Rotation of **1** in the active site of III mutant.

**Description:** At first, testosterone (**1**) is bound in pose 15, perpendicularly with respect the haem plane and interacting with T260 and A330 through its hydroxyl and carbonyl moieties, respectively. Gradually, **1** assumes a lying pose over its α-side, and the subsequent (**1**)CO-T327 and (**1**)OH-A87 interactions start driving a counterclockwise rotation of **1** inside the haem pocket. Afterwards, (**1**)CO-G265 followed by (**1**)OH-T327 interacions complete the 180º rotation of **1**, which further on assumes its final binding pose 2 perpendicular to the haem plane.

**Supplementary Video 2:** Binding trajectory of **1** in I-- mutant using accelerated MD simulations.

**Description:** Fetching of (**1**) takes places through the F-G loop / β1 hairpin. The substrate enters in the enzyme pocket from the access channel 2a and there it reorients until a network of coupled conformational changes that allows the path of (**1**) towards the active site occur simultaneously. The G helix adopts a bend conformation, which impacts F helix and β1 sheet conformation, and in turn shifts B' helix and retreats β4 sheet, enabling **1** progression to the catalytic centre.
